# From Data to Theory: Understanding the Role of Naive Learners in Cultural Change

**DOI:** 10.1101/2022.05.20.492808

**Authors:** Michael Chimento, Lucy M. Aplin

## Abstract

A change to a population’s social network is a change to the substrate of cultural transmission, affecting behavioral diversity and adaptive cultural evolution. While features of network structure such as population size and density have been well studied, less is understood about the influence of social processes such as population turnover— or the repeated replacement of individuals. Experimental data has led to the hypothesis that naive learners can drive cultural evolution by being better samplers, although this hypothesis has only been expressed verbally. We conduct a formal exploration of this hypothesis using a generative model that concurrently simulates its two key ingredients: social transmission and reinforcement learning. We explore how variation in turnover influences changes in the distributions of cultural behaviors over long and short time-scales. We simulate competition between a high and low reward behavior, while varying turnover magnitude and tempo. We find optimal turnover regimes that amplify the production of higher reward behaviors. We also find that these optimal regimes result in a new population composition, where fewer agents which know both behaviors, and more agents know only the high reward behavior. These two effects depend on network size, density, behavioral transmissibility, and characteristics of the learners. Our model provides formal theoretical support for, and predictions about, the hypothesis that naive learners can shape cultural change through their enhanced sampling ability, identified by previous experimental studies. By moving from experimental data to theory, we illuminate an under-discussed generative process arising from an interaction between social dynamics and learning that can lead to changes in cultural behavior.

## Introduction

Culture, broadly defined as shared socially-learned traits that persist over generations, is widespread across taxonomic groups including birds, cetaceans and primates (Aplin, 2019; Rendell and Whitehead, 2001; Schuppli and van Schaik, 2019; Whitehead et al., 2004; Whiten, 2019, 2021). Variants of cultural traits are inherited through social learning and compete with each other, potentially changing over time to become more adaptive in a process of cultural evolution (Creanza et al., 2017*a*; Mesoudi, 2017; Mesoudi and Thornton, 2018; Mesoudi et al., 2004; Whiten et al., 2011). Critical to the adaptiveness of culture are processes that increase the frequency of higher reward behaviors. Transmission biases (Boyd and Richerson, 1988; Kendal et al., 2018; Laland, 2004; Rendell et al., 2011), teaching, and meta-cognitive abilities (Dean et al., 2012; Heyes, 2016, 2018; Legare, 2017; Tennie et al., 2009) have all been suggested as mechanisms that enable the adaptiveness of culture. However, simpler mechanisms such as the introduction of naive learners via population turnover, and reinforcement learning that occurs during the maintenance of socially acquired behavior (Galef Jr, 1995) (called guided variation by Boyd and Richerson (1988)) could also increase adaptiveness. Crucially, these mechanisms are present as part of the phenotypes of many social species (Boyd and Richerson, 1988; Chimento et al., 2021, 2022; Sasaki and Biro, 2017; Warner, 1988). In our recent experiment, we found that new learners preferentially adopted a more efficient foraging behavior rather than a less efficient established tradition, supporting the hypothesis that new learners can drive cultural evolution by better sampling the available behavior space without interference from prior experience (Chimento et al., 2021). However, this was a verbal hypothesis drawn from limited experimental data. In this study, we use a mechanistic computational model to explore turnover theoretically and provide robust predictions under minimal assumptions.

Culture is influenced by social structure and processes, with social network dynamics having a profound effect on cultural evolution (Nunn et al., 2009; Whitehead and Lusseau, 2012). Recently, effects of demographic structure, immigration, and population growth have all received increased attention. In age-structured populations, different social learning strategies may evolve, such as copy-the-young or copy-the-old, depending on environmental variability (Deffner and McElreath, 2022). In evolutionary linguistics, social processes, such as immigration and population growth, have been associated with the reduction of morphological diversity (Dale and Lupyan, 2012; Lupyan and Dale, 2010; Raviv et al., 2019; Wray and Grace, 2007), with new learners exerting selection pressure for simpler, more structured languages (Kirby et al., 2014). In the context of animal culture, the introduction of new individuals has been shown to improve flight routes, spawning sites, and foraging behaviors (Chimento et al., 2021; Sasaki and Biro, 2017; Warner, 1988). Despite the hypothesized importance of new learners, population turnover, a fundamental social process that can introduce new learners, remains one of the most understudied social processes in the field. We note that our use of “population turnover” differs from the introduction of novel traits (Acerbi and Bentley, 2014; Bentley et al., 2007).

Turnover, a gradual replacement of individuals in a population, is a common feature of nearly all animal social systems, often caused by fission-fusion movements, or dispersal between groups, or generational change (Aureli et al., 2008; Silk et al., 2014; Sueur et al., 2011). Turnover is a means by which new learners are introduced into a population, and culture depends on transmission to new learners. However, turnover has rarely been the object of study itself. Cultural diffusion experiments typically introduce naive learners, yet the rate or magnitude of replacement is not varied (Mesoudi and Whiten, 2008; Whiten and Mesoudi, 2008). Similarly, prior computational models of culture have included turnover, but it was not the object of study and occurred in the same manner for all conditions (Kirby and Tamariz, 2022; Smith, 2002). The lineage of classic differential equation models of cultural diffusion, “fundamental diffusion models”, such as the mixed-influence model (Bass, 1969), assume a full mixing population (complete network), a fixed number of potential adopters, and a binary diffusion process (individuals do or don’t adopt a behavior). A prior model of guided variation by Boyd and Richerson also assumed a full mixing population, and turnover occurred for all individuals only at the end of each generation (Boyd and Richerson, 1988). Neither of these formal models could explicitly specify and explore competition between behaviors in structured, dynamic networks. Given the ubiquity of turnover in social species and its central role in cultural evolution, it is crucial to further our understanding of its influence on socially learning species beyond the human behavioral context.

There have been only 3 empirical studies that have explicitly investigated the effect of turnover on animal cultures. First, after an experiment in Blue-headed wrasse (*Thalassoma bifasciatum*) that entirely replaced sub-populations with naive individuals, groups chose more suitable spawning sites (Warner, 1988). These newly chosen sites were chosen again after a second replacement event (Warner, 1990), suggesting that the original culturally inherited preferences were the suboptimal result of cultural inertia. Another study found that successive replacement of pairs of homing pigeons (*Columba livia*) in a transmission chain increased route efficiency (Sasaki and Biro, 2017). Finally, our recent study directly compared foraging cultures in static and turnover micro-populations of captive great tits (*Parus major*), showing that turnover populations were much more likely to shift to a more efficient foraging behavior over an established, inefficient tradition (Chimento et al., 2021). These studies hypothesized that the improvement of cultural adaptiveness in groups with turnover was due to naive individuals introduced new innovations (Sasaki and Biro, 2017), or improved the sampling of behavior space by naive individuals (Chimento et al., 2021; Warner, 1988, 1990). However, these were proposed hypotheses drawn from experimental data. To our knowledge, no theoretical work exists that explores the mechanism by which naive individuals introduced via turnover may improve cultural adaptiveness of groups. Prior models have only asked adjacent questions in the context of cultural diversity, rather than cultural adaptation, exploring population contact between already knowledgeable individuals, rather than naive individuals (Creanza et al., 2017*b*; Somveille et al., 2018).

In order to better understand how turnover might affect cultural evolution, we used a computational model to explore the effect of turnover in structured populations, where turnover was designed to mimic immigration and emigration between a focal population and a source of naive individuals. We simulated the diffusion of a novel, higher reward behavior, which competed against an established, low reward behavior, based on the design of our previous experiment (Chimento et al., 2021). We varied the intensity of turnover by changing 1) the magnitude, or what percentage of the population was replaced, and 2) the tempo, or how frequently replacement events occurred. We further varied network size, density, the transmissibility of behaviors, as well as the rules that agents used to make behavioral decisions, with the expectation that the effect of turnover might depend on these variables. Our results showed that as turnover changed the composition of repertoires in the population and agents’ beliefs about their own behavioral choices in predictable ways. When compared to static populations, these changes could lead to an increased relative proportion of high-reward behavior when turnover was low intensity, or behavioral extirpation when turnover was high intensity.

## Methods

Our model simulated dynamic social networks of size *N*, populated by reinforcement learning agents (Sutton et al., 1998), whose behavioral repertoire and behavioral preferences could change over time. Agents present in the first generation were initialized with knowledge of a low-reward behavior *a*, which was intended to represent an established tradition. One agent was randomly chosen as a “seed” agent, with knowledge of a high-reward behavior *b*. The simulation’s design mirrored our prior experiment using great tits (Chimento et al., 2021), where a lower reward tradition was allowed to spread through groups, and then groups were either held static or had 2 of 6 members replaced by naive individuals every week. To reduce variance, we began simulations after one individual had learned the higher-reward behavior. Figure 1A illustrates initial conditions and a sample simulation.

**Figure 1:**
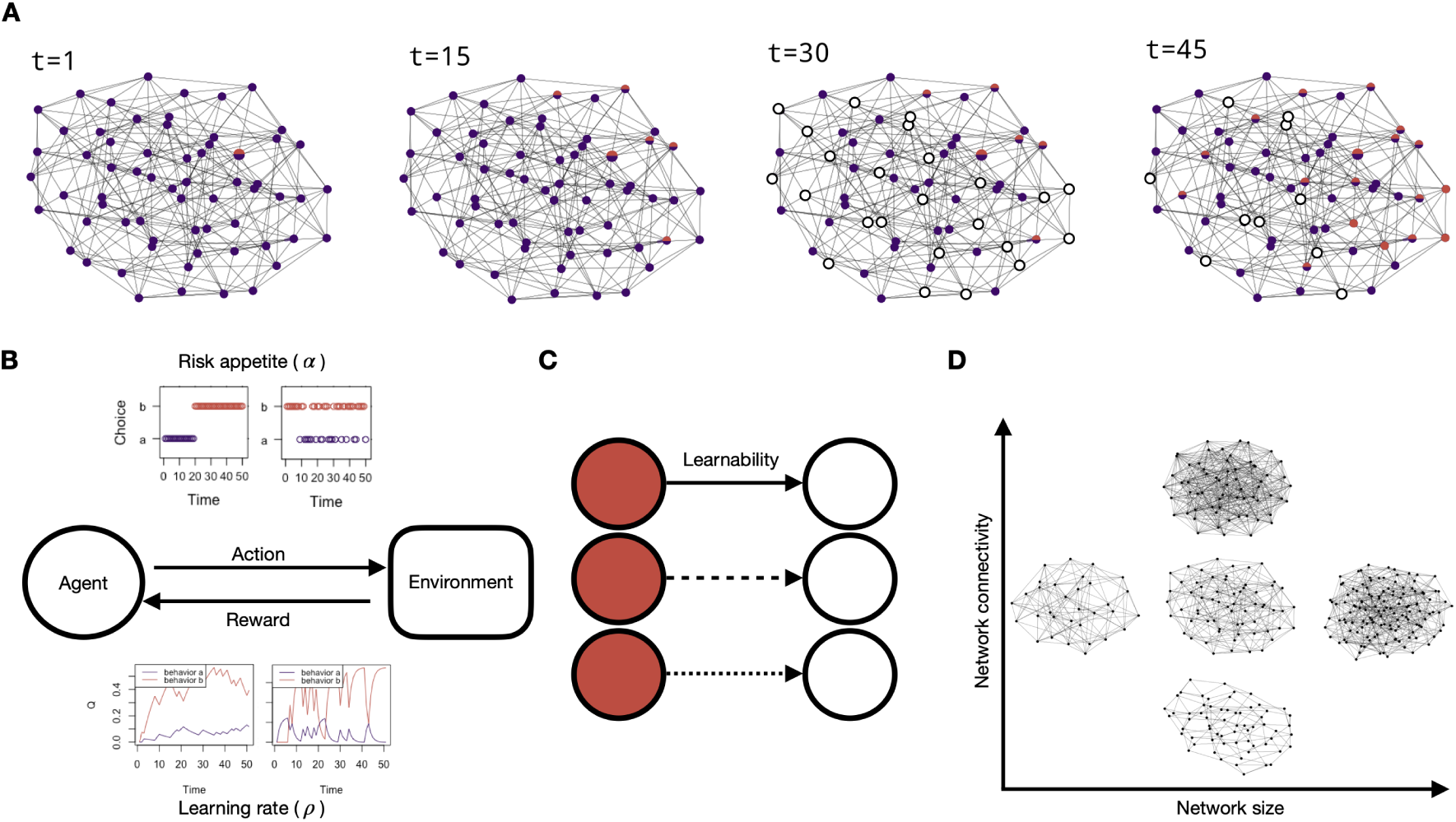
Model summary. **A)** Example of a simulation’s progression at the reference parameter constellation. At timestep *t* = 1, all agents held the low-reward behavior (*a*, purple) in repertoire. One seed agent held an alternative high-reward behavior (*b*, red). At *t* = 15, *b* spread to several agents via social transmission. These agents may now produce either *a* or *b* each timestep. A turnover event occurred at *t* = 30. Turnover magnitude is *T_m_* = *N ∗ .*375, thus 24 agents are replaced by naive agents with knowledge of neither behavior (white). At *t* = 45, some naive agents have now acquired knowledge of one behavior or the other. We varied parameters that defined **B)** the choice and reward functions in the sub-model of production, **C** the transmissibility of behaviors, and **D)** network size and connectivity.

In each time-step, agents could possibly acquire knowledge about *b*, and also chose one behavior from their repertoire to produce, for which they received a reward. Agents’ learning dynamics for both acquisition and production were defined by a simplified implementation of our prior model of cultural diffusion (Chimento et al., 2022). Behaviors were added to an agent’s repertoire through a network-based diffusion analysis dynamic (Franz and Nunn, 2009; Hasen-jager et al., 2020; Hoppitt and Laland, 2013), which was agnostic to the mechanism responsible for social transmission, and could represent anything from enhancement to imitation (Hoppitt and Laland, 2013). Agents could acquire knowledge of behaviors either through a very low probability of asocial innovation, or observation of neighboring agents, defined by a social network. Acquisition was thus conditioned on both association and production: the more productions of a novel behavior an agent observed, the more likely it was to acquire knowledge of that behavior.

Agents used a Rescorla-Wagner (RW) model to select a behavior for a given timestep (Sutton et al., 1998). This model made agents reinforcement learners who chose behaviors from their repertoire based on previously received rewards. Over time, agents’ expected values about behaviors updated with experience, influencing the probabilities of choosing behaviors from their repertoire.

We varied the intensity of turnover the population would experience by either leaving populations as static, or by replacing a certain number of agents (turnover magnitude, *T_m_*) at a set interval (turnover tempo, *T_t_*). Each combination of these two parameters defined a turnover regime. Naive agents replaced older agents with a probability proportional to their time spent in the population and inherited the network connections of their predecessors. They had to learn behavioral knowledge by observing the productions of neighbors since they had no prior knowledge of *a* or *b*. We also varied parameters related network size and connectivity, behavioral transmissibility, and reinforcement learning to understand how these might interact with turnover (summarized in figure 1B,C,D).

In order to analyze the model, we conducted a sensitivity analysis where each parameter of the model was varied in turn while all others were held constant. We conducted 100 simulations each lasting 10, 000 timesteps for each parameter constellation. In each simulation, we recorded the distributions of repertoire states, behavioral productions at each timestep and expected values of behaviors. We then compared these key measurements to static populations in order to assess the relative effect of each turnover regime, as well as the how the effect of turnover interacted with other parameters in the model.

The equations of both sub-models are defined below.

### Sub-model of acquisition

Each time-step, an agent *i* could independently innovate or socially learn behavior *a* or *b* from their neighbors, as defined by the social network. This probability of acquisition was defined by Equation 1, where it was influenced by a base learning rate (*λ*) and a transmission function *T* that accounted for the associates of *i* defined by the social network (binary vector *a_i_* where *a_i_*_,*j*_ = 1 if associated, else 0) and their behavioral productions (*w_k_*(*t*)) where *k ∈ {a*, *b}*.

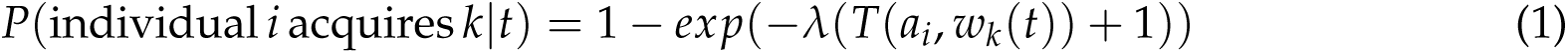

The +1 accounts for asocial innovation of the behavior by an agent. We were primarily interested in cultural behaviors spread by social transmission, so we set the probability of independent innovation to a very low value for both behaviors (*λ* = 0.001, resulting in innovation probability of .00099 per time-step). The transmission function *T* (Equation 2) contained the social transmission rate *s*, which defined how easily an agent might socially acquire knowledge of a behavior (transmissibility). This multiplied the sum of associates’ behavioral productions, defined by transmission weight *w_k_*_,*j*_(*t*).

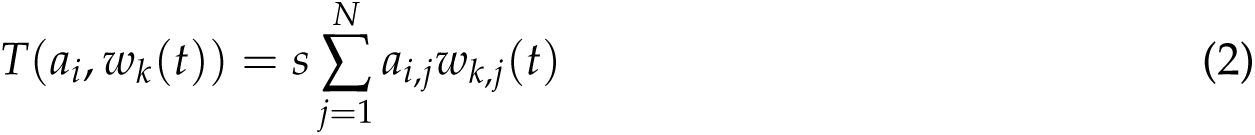

The transmission weight (*w_k_*_,*j*_(*t*)) was a value of either [0, 1], depending on whether individual *j* produced behavior *k* in timestep *t*. If agent *j* produced behavior *a* in timestep *t*, *n_a_*_,*j*_(*t*) = 1 and *n_b_*_,*j*_(*t*) = 0. The transmission weight was an important connection between the sub-model of acquisition and the sub-model of behavioral production. The probability of acquisition depended on who an agent was neighbors with, and what those neighbors produced.

### Sub-model of behavioral production

Our sub-model of behavioral production was a simple reinforcement learning model with two parameters: learning rate and risk-appetite. In each time-step, agents produced one behavior *k* from their repertoire using the sub-model of production, for which they received a reward. Agents received deterministic rewards for producing either behavior defined as

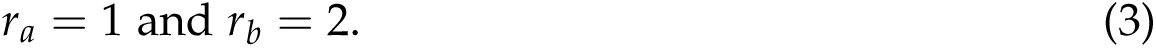

After producing a behavior, agents updated their expected value for behavior *k* using *Q*(*k*):

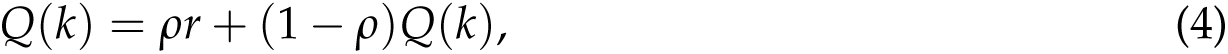

where *ρ* was a learning rate parameter that controlled how strongly the received reward influenced the expected value *Q*. The un-chosen behavior *j* was updated as

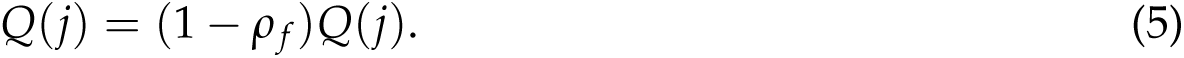

When the forgetting rate *ρ _f_* = 0, agents never lost information about previous received rewards. When *ρ _f_ >* 0, agents forgot the value of un-chosen options over time.

In the first time-step of a simulation, all agents in a population were assumed to be knowledgeable of the low reward behavior, yet held no expectations about its value *Q*(*a*) = 0. One agent *j* was seeded with knowledge about high-reward behavior *b* to eliminate noise at the beginning of simulations due to stochastic differences in the time before the initial innovation event. This seed agent held *Q*(*b*) = 0, giving it an equal chance of producing either behavior. In turnover conditions, naive agents entered populations without knowledge of either behavior. Upon acquisition of behavior *k*, *Q*(*k*) = 0, conservatively assuming that agents did not attend to rewards before acquisition, and held no prior expectations about the reward.

Using a softmax function, expected values were transformed into the probability of producing a behavior. *α* determined the risk-appetite of the agents. If *α <* 1, agents were risk-averse, and would prefer to produce the behavior for which they held the highest expected value. If *α >* 1, agents were risk-tolerant, and would engage in exploratory sampling of behavior, insensitive to expected values. When *α* = 1, agents were linearly sensitive to differences in expected values.

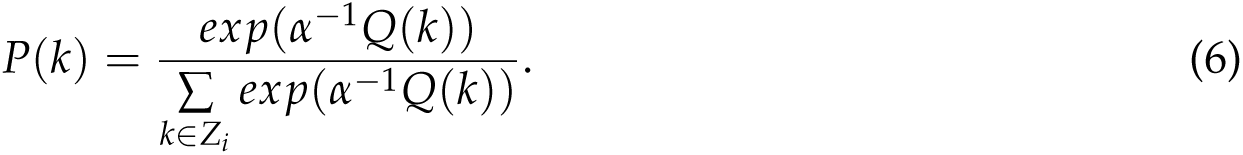

When agents knew only 1 behavior *k*, *P*(*k*) = 1.

### Measurements and conditions

We manipulated the magnitude of turnover *T_m_* from 0% of the population to 100% of the population in increments of 12.5%. We also varied the tempo of turnover *T_t_* from 10 to 100 in increments of 10. We conducted an additional exploration of turnover tempos beyond these values, summarized in figure S1.

We tested random regular networks of size *N* agents. In random regular networks, agents are randomly connected to *k* other agents. We chose random regular architecture because network degree was important to standardize, as it directly influenced the probabilities calculated in the sub-model of acquisition (Equation 2). We tested 3 populations sizes (*N ∈ {*32, 64, 128*}* along with 3 levels of network connectivity by changing the relative proportion of *k* against *N* (*k ∈ {*0.0625*N*, .125*N*, .25*N}*). For example, a network with *N* = 64 and *k* = 0.125*N* yielded *k* = 8—comparable to a network where *N* = 32 and *k* = 0.25*N*, and *N* = 128 and *k* = 0.0625*N*. This allowed us to compare network sizes while controlling for degree, and vice-versa.

Within the sub-model of acquisition, we varied the transmissibility of behaviors, or how easily an agent could socially acquire either the novel or established behavior (*s ∈ {*2, 5, 8*}*). Within the sub-model of production, we varied decision-making rules. We tested 3 different parameterizations of the learning rate parameter (*ρ ∈ {*.01, 0.1, 0.99*}*); and risk-appetite parameter (*α ∈ {*0.5, 1.0, 2.0*}*). In the main text, we leave *ρ _f_* = 0 for simplicity, so there is no forgetting of expected value. In the supplementary, to compare the effect of information loss from memory and from turnover, we provide an additional analysis of the forgetting rate *ρ _f_* from .1 to .9 in increments of .1.

In order to understand the effect of turnover on cultural evolution, we conducted a sensitivity analysis in which we varied one parameter at a time, while holding all others at the reference setting. Our reference setting was

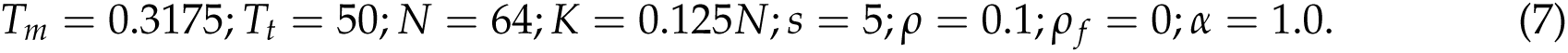

For each point in parameter space described above, we recorded 100 simulations, each running for 5, 000 time-steps. At each time-step, we recorded behavioral frequencies, agents’ repertoires, and agents’ expected values (*Q*(*k*)) about the behavior in their repertoires.

## Results

Our simulation model aimed to formally explore the hypothesis that the introduction of naive learners can drive cultural change to more or less efficient cultural traits. The simulations mirrored our prior design which experimentally explored this question in great tits using captive micro-populations (Chimento et al., 2021), but extended our exploration far beyond what was possible in that experiment. Populations began with knowledge of a single, low-reward behavior, but had the opportunity to learn about and use a higher-reward behavior. We measured behavioral productions, repertoire states, and expected values of each behavior, all while varying the magnitude and tempo of turnover. Our simulation model allowed us to compare the effects of different turnover regimes across various population sizes, network connectivities, and learning rules.

We first explored static populations as a baseline for comparing all other turnover regimes. In the reference static condition (Equation 7), the novel behavior spread to all individuals in a population within a mean*±*SD of 318 *±* 47.2 time-steps. Once this diffusion finished, the population was composed only of polymorphic agents who knew both behaviors (figure 2,A). Populations switched their majority preference from *a* to *b* by timestep 189, on average. Over the entire simulation, the high-reward behavior represented a mean proportion of 0.709 *±* 0.121 of all behaviors produced in the static reference constellation (figure 2,C). This value was similar across other parameter values (range: 0.691 *−* 0.717), with the exception of risk appetite. Risk averse populations obtained a mean proportion of 0.852 *±* 0.142, while risk tolerant populations obtained 0.604 *±* 0.108. This was due to agents’ relatively stronger or weaker preference for higher-valued behavior.

**Figure 2:**
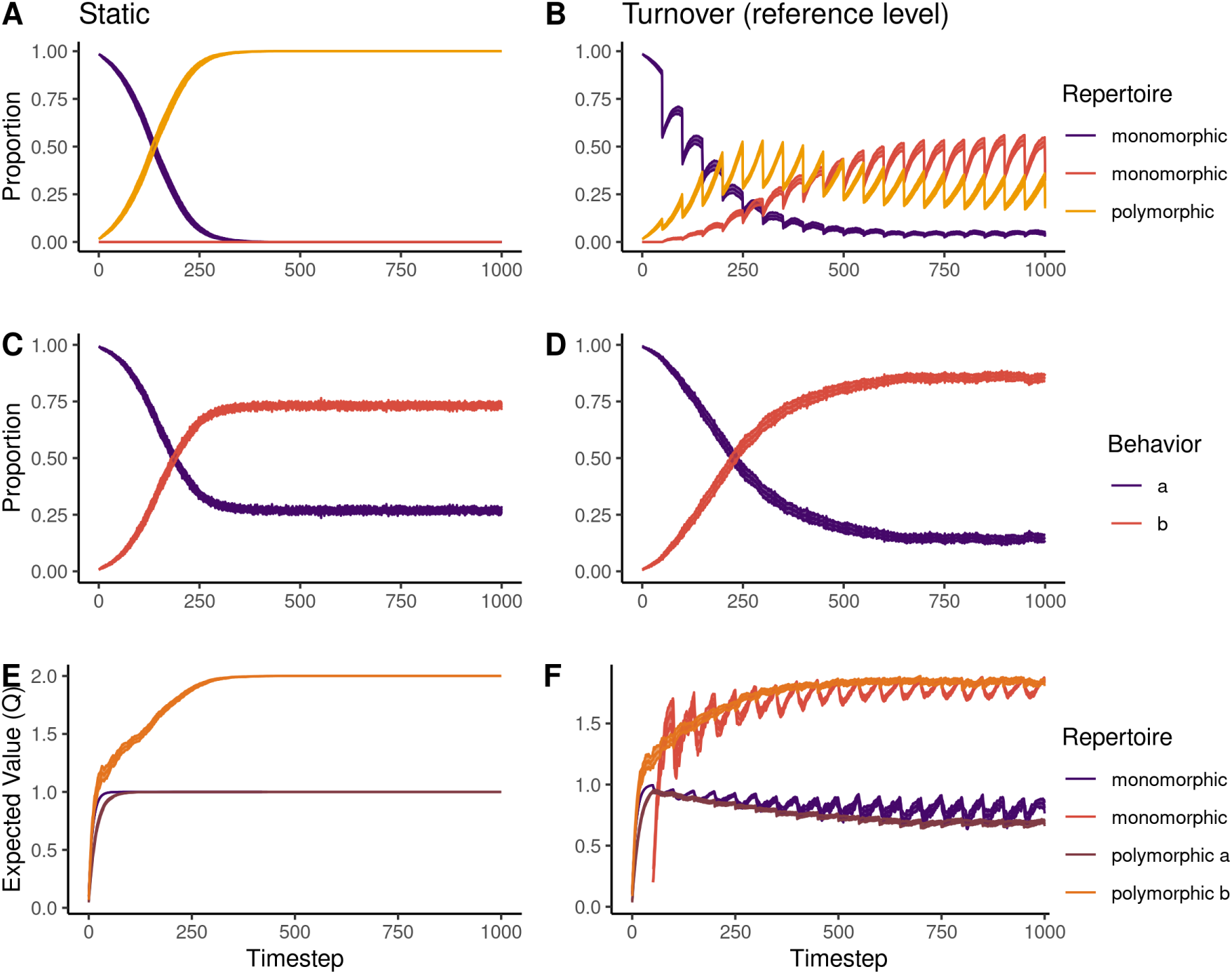
Temporal simulation dynamics. Mean and bootstrapped 95% CL of 3 key metrics (rows) in static and turnover reference level simulations over the first 1000 timesteps (dynamics maintain a steady state for the remaining timesteps.) The first row is the proportion of individuals in each repertoire state. Turnover populations experience an rise in polymorphic agents, followed by a steady rise in high-reward monomorphic agents. The second row shows the relative proportion of each behavior that was produced each timestep. The high-reward behavior *b* represents a higher proportion of observed behaviors compared to static populations. The third row shows the expected value that knowledgeable agents held about either behavior conditional on repertoire state.

In the static reference constellation, agents’ mean expected values were *Q_a_* = 0.998 *±* 0.029 and *Q_b_* = 1.972 *±* 0.146 (figure 2,E). These were slightly lower than the true rewards due to time spent learning the rewards at the beginning of the simulation (Equation 4).Expected values were largely similar across parameter values with the exception of learning rate (*ρ*), where a slow learning rate resulted in *Q_b_* = 1.86 *±* 0.391 and a fast learning rate resulted in *Q_b_* = 2.00 *±* 0.056.

Thus, the two factors that affected outcomes between static populations were 1) risk-appetite, in that risk-averse agents obtained slower diffusions, but disproportionately shifted probability to choosing the high-reward behavior, and 2) learning rate, in that agents with slower learning rates took longer to learn the full value of behaviors. We expected these two parameters to also play an important role in the effect of turnover, as more frequently produced behaviors would be more likely to be transmitted, and agents would also be time limited in how long they could learn value.

### The effect of turnover magnitude and tempo

Next, we measured how population turnover influenced simulation dynamics. At the reference constellation, we found that the population underwent a gradual recomposition of repertoires (figure 2,B). The number of low-reward monomorphic agents steadily declined over time, obtaining a mean proportion of 0.067 *±* 0.131. There was an initial increase and subsequent decline in polymorphic agents, with a mean proportion of 0.246 *±* 0.101. Polymorphic agents eventually became outnumbered by high-reward monomorphic agents who comprised nearly half of the population (0.448 *±* 0.155). These relative proportions of each repertoire stabilized after approximately 700 time-steps.

There was a slower adoption of the high-reward behavior (majority preference by timestep 227), with static populations obtaining a higher relative efficiency in the early stages of the simulation. However, in the long run, *b* made up a higher proportion of overall behaviors produced by the population compared to static (mean SD proportion 0.831 *±* 0.163, figure 2,D). Turnover populations held slightly lower expected values for both behaviors (mean SD *Q_a_* = 0.705 *±* 0.098, and *Q_b_* = 1.782 *±* 0.138. The expected values by repertoire state showed evidence that polymorphic individuals largely acquired the high-reward behavior first, and the low-reward behavior second. Monomorphic agents (figure 2F) held lower expectations about the high reward behavior compared to polymorphic individuals, and polymorphic individuals held lower expectations about the low-reward behavior compared to monomorphic individuals. This difference was caused by the time taken for high-reward monomorphic individuals to acquire the low-reward behavior (all values summarized in table S3).

Our results showed that turnover can improve the relative ”efficiency” of the cultural state of the population through a process of information loss and recomposition, while simultaneously reducing the carrying capacity for cultural traits. However, the gradual change in repertoires and slower adoption of the high-reward behavior did not quite match our prior experimental results, which showed a relatively fast switch to the high-reward behavior in populations that experience turnover (Chimento et al., 2021). We return to the reasons for those differences in results section “Turnover could benefit populations in the short term”.

### Optimal, neutral and sub-optimal regimes

We then explored how altering turnover magnitude and tempo influenced simulation dynamics (detailed summary statistics in Table S1 and Table S2). Holding turnover tempo at *T_t_*= 50, we found that the mean proportion of high reward behavioral productions increased as magnitude increased, although this relationship was quadratic with a maximum when turnover magnitude *T_m_* = .3175 (figure 3A, rows). While holding magnitude constant at *T_m_* = .3175, we observed a similar pattern of increasing proportion of high-reward behaviors as tempo increased, with a maximum at *T_t_* = 50 (figure 3A, columns). At slower tempos, populations could experience higher magnitudes with no detrimental effect. We extended our exploration of tempo up to 2048 timestep intervals, and found that as tempo decreased, the effect of turnover magnitude lessened, until behavioral frequencies were little changed from static populations (figure S1).

**Figure 3:**
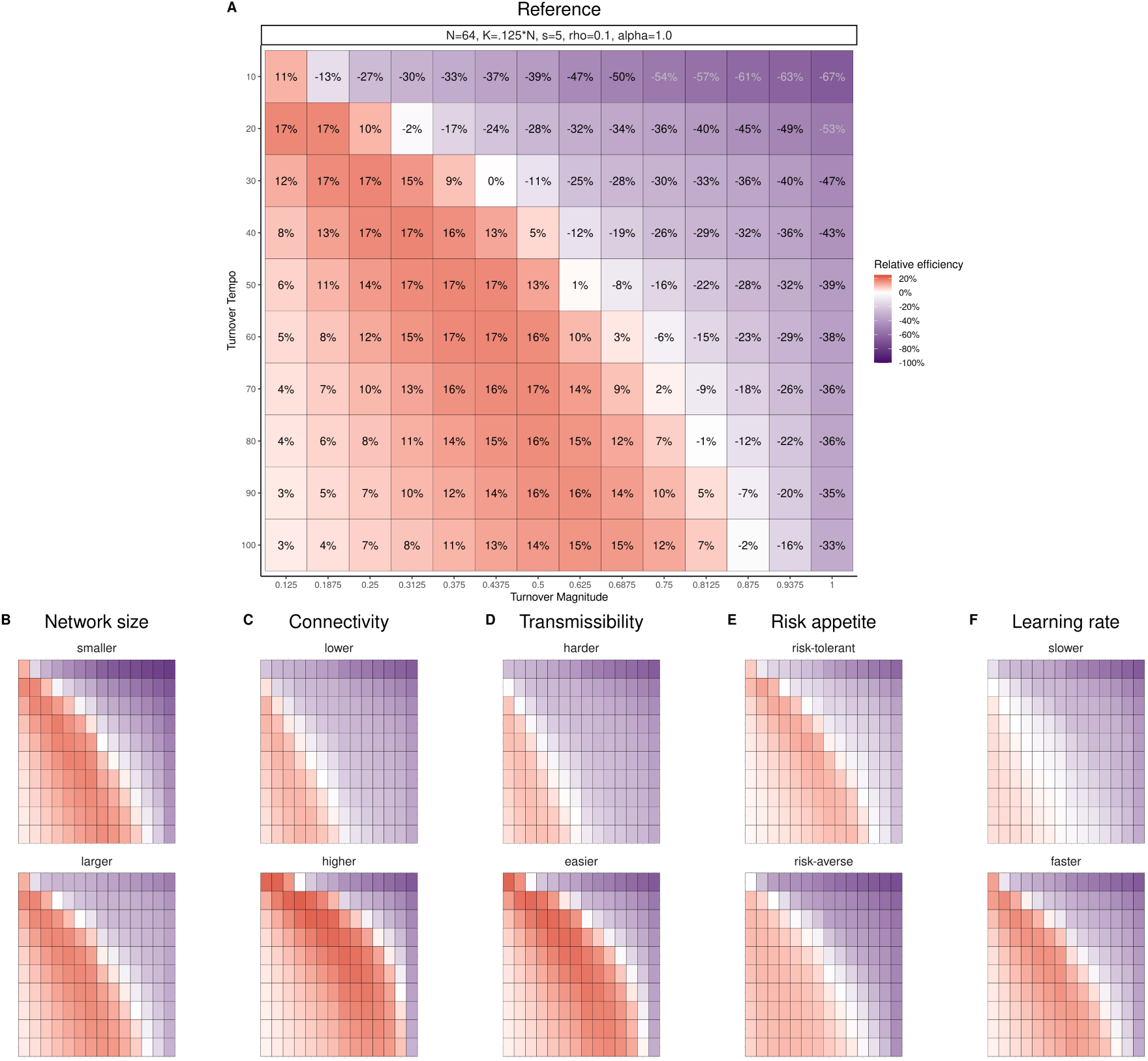
Optimal turnover regimes depended on network size, density, and transmissibility of behavior. Mean relative efficiency (fill, label) shown for 100 simulations under each turnover regime: the combination of turnover magnitude (x-axis) and turnover tempo (y-axis). Relative efficiency was measured against the proportion of *b* produced by the static populations under a given condition, where the color white is equivalent performance, salmon is over-performance and purple is under-performance. Smaller, sparser networks, and cases where behaviors are not easily transmissible were less tolerant of turnover, yielded less benefit from optimal regimes and under-performed under a larger number of regimes. The opposite was true of larger, denser networks with more transmissible behaviors.

In summary, regimes with slow tempos and low magnitudes resulted in equivalent relative efficiency to static populations. Relative efficiency of regimes increased as magnitude and tempo increased to a certain threshold, with faster tempos required when magnitude was small, and larger magnitudes when tempo was slow. When turnover increased in intensity beyond this point, the relative efficiency decreased and the probability of behavioral extinction increased. Few individuals held either behavior in repertoire, as turnover events removed too many knowledgeable individuals or occurred so frequently that agents were not present in the population for long enough to acquire either behavior. The relative frequencies of either behavior oscillated around .5, since similar numbers of individuals knew either behavior, but not both. Regimes with large magnitudes and fast tempos resulted in very poor performance, especially when tempo was fastest.

### The effect of turnover depended on network structure, transmissibility and production rules

The thresholds for the transitions between neutral, optimal and sub-optimal relative efficiency depended on network size, connectivity and transmissiblity (figure 3B, C, D). Detailed summary statistics for these constellations at the reference level are presented in Table S4. In summary, strong turnover regimes were most detrimental in small networks, sparse networks, and for behaviors which were difficult to socially learn. Large networks, dense networks and easily learnable behaviors all increased the number of turnover regimes that a population could undergo without negative performance compared to static populations. Additionally, the benefits of optimal regimes improved under these same conditions. However, with the exception of network size, these conditions also increased the tempos required to out-perform static populations.

The effects of network parameters were caused by how they altered the probabilities of behavioral extinction and transmission. Network size determined the amount of memory available within a population. Altering the size changed the probability of behavioral extinction, with a larger risk of extinction (and reduced probability of any innovation) in smaller networks. Altering network density indirectly increased the transmissibility of behavior. The more associates any agent had, the more likely they were to observe either behavior, which increased the rate of social transmission. Of course, directly increasing the transmissibility of the behavior via the social learning rate also hastened acquisition by naive agents (NB: this parameter did not increase the probability of independent innovation). The similar effect of density and transmissibility are highlighted in how similarly the re-composition of the population evolved when *K* = .25*N* or *s* = 8 (figure S3). Regardless of the mechanism, if behaviors were more easily acquired, populations were more resilient against effects of extreme turnover regimes.

Finally, the flexibility of our production sub-model allowed us to test the effect of turnover on different types of learners. Populations composed of risk-tolerant agents saw less benefit from turnover, as polymorphic individuals were more likely to sample either behavior irrespective of their relative expected value (figure 3E). Interestingly, the increase in relative efficiency in risk-averse populations was muted, with a maximum at about 13%. Risk-averse populations obtained higher proportions of high-reward monomorphic agents (mean 0.390 *±* 0.130 compared to reference level 0.278 *±* 0.117), and polymorphic agents were more likely to choose the high-reward behavior. However, the consequence of increasing the sensitivity to expected values meant that static populations also produced more of behavior *b* over a longer period of time. This indicated that our parameterization of behavioral conservatism didn’t capture the effect seen in our experiment, which we return to in the final section of the results. Populations with a very slow learning rate (*ρ* = 0.01) saw the least benefit from turnover (figure 3F). This was because agents did not last long enough in the population to learn the differences in values of each behavior (mean *Q_a_* = 0.062 *±* 0.011, *Q_b_* = 0.206 *±* .031), making *a* relatively competitive against *b*. As a consequence, slow learning rate also obtained a higher relative proportion of monomorphic learners compared to the reference level (Table S4).

### The effect of turnover on population composition

In order to understand the mechanisms that mediated the relationship between turnover and cultural state, we explored the knowledge states of agents in terms of their repertoire and expected values. In static populations, 100% of agents became behaviorally polymorphic (knew both behaviors). As magnitude or tempo increased towards their optimal values, the proportion of high-reward monomorphic agents (only knew behavior *b*) increased, with a maximum of .594 [0.469, 0.719] at optimal values of magnitude (shown in figure 4A) and tempo. The proportion of polymorphic and of low-reward monomorphic agents (only knew behavior *a*) remained lower than high-reward monomorphic agents. As turnover magnitude and tempo reached more extreme levels, the proportion of high-reward monomorphic agents fell. Interestingly, the proportion of low-reward monomorphic agents began to rise to match the curve of high-reward monomorphic agents (figure 4A). This was caused by 1) the rare independent innovation and subsequent diffusion of the low-reward behavior to completely naive agents, and 2) the properties of reinforcement learning over short time periods. Under strong turnover regimes, the two behaviors became more competitive against each other because knowledgeable agents spent less time in the population and had less experience with either behavior. They therefore weren’t able to fully learn their value.

**Figure 4:**
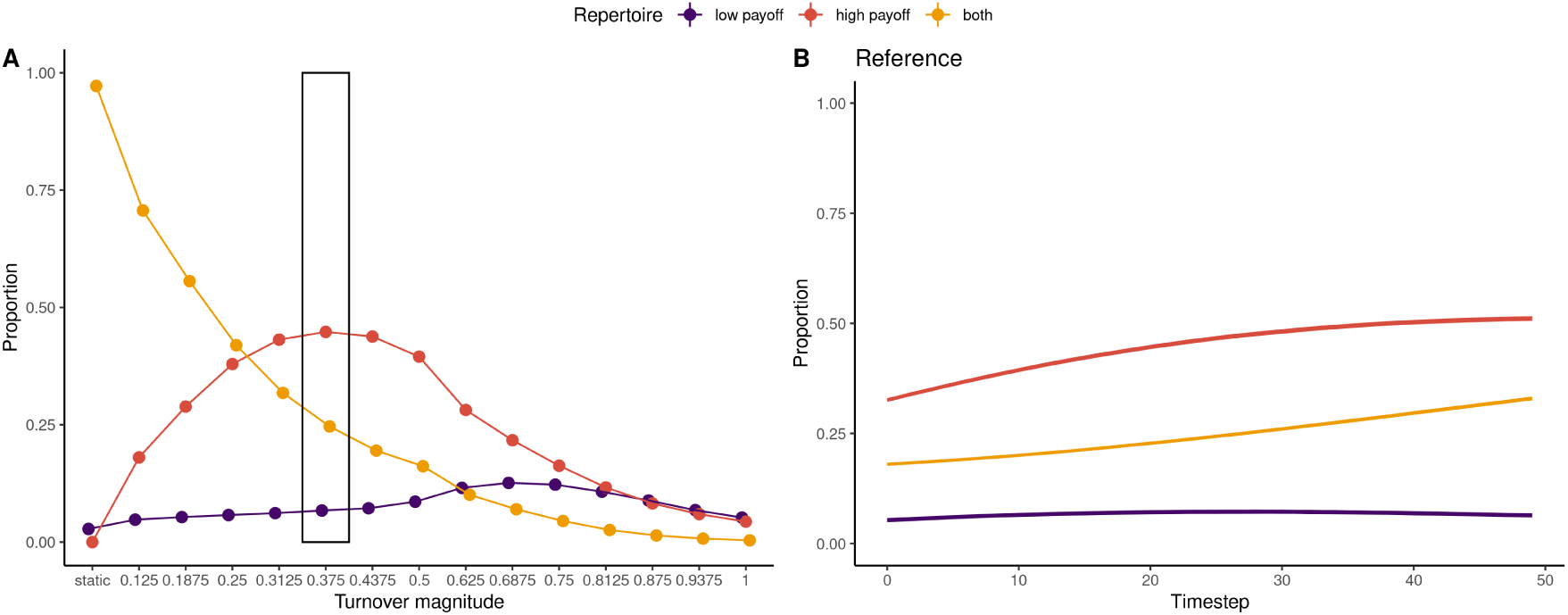
Turnover acts on the composition of cultural repertoire. **A)** Comparison of mean and bootstrapped 95% CL of the proportion of agents that were knowledgeable of *a* only (purple), *b* only (salmon), and both behaviors (yellow). As turnover magnitude increased, the number of polymorphic individuals decreased. The optimal magnitude for the reference tempo (rectangle), and vice versa, occur when the proportion of high-reward monomorphic agents exceed the number of polymorphic agents. **B)** Temporal dynamics of population composition (mean, bootstrapped 95% CL) in timesteps following turnover event in reference constellation. After turnover, the number of high-reward monomorphic agents rose faster than the number of polymorphic individuals.

Given the importance of information loss for the effect of turnover, we predicted it was possible that a forgetting rate (*ρ _f_*) might induce the same changes to relative efficiency in the absence of turnover. We conducted a sensitivity analysis of varying levels of *ρ* and *ρ _f_* in static populations, and found that increasing *ρ _f_* could increase relative efficiency when compared to *ρ _f_* = 0 (figure S2) when the the learning rate was also relatively high. Otherwise, it led to poorer performance. While the analogy between forgetting rates and turnover is intriguing, the mechanisms that change the distribution of cultural traits are completely different than the turnover model. A forgetting rate did not capture the changes to population composition that occurred when individuals were replaced.

Namely, a model of static populations with a forgetting rate did not capture the temporal dynamics of recomposition. We found that after a turnover event, populations’ composition changed in a consistent manner conditional on the parameter constellation (e.g. figure 4B). The effects of turnover ultimately depended on the probabilities of naive individuals to transition from naive to behaviorally monomorphic or polymorphic. All parameters influenced recomposition following turnover events to some degree, and have visualized these differences in figure S3. We previously hypothesized that turnover might have an especially strong effect in behaviorally conservative species (Chimento et al., 2021). After each turnover event, risk-averse populations obtained a much sharper rise in high-reward monomorphic agents compared to reference level, supporting our hypothesis. However, as risk-appetite also caused static populations to produce the high-reward behavior more frequently, negating any benefits from recomposition. In the next section, we summarize how an alternative parameterization of behavioral conservatism generated data that better fit our previous experimental results.

### Turnover could benefit populations in the short term

The inspiration for this model was the verbal hypothesis inspired by our prior experiment with great tits, where we found that turnover provided benefits to populations in the short term. Turnover populations switched over to a high-reward alternative faster than static populations after a handful of turnover events Chimento et al. (2021). In our model, turnover populations did not switch over before static populations, and only outperformed them in the long run through population recomposition. However, there were two key changes we predicted would generate data that more closely matched our prior results and verbal hypothesis. These modifications are detailed in appendix A.

We found that the risk-appetite parameter alone did not greatly inhibit static populations from switching over to behavior *b*. To change this, we introduced a choice autocorrelation parameter, which is a widely used parameter in reinforcement learning models in humans (Katahira, 2015). When this parameter is above zero, agents are more likely to repeat recent choices, further slowing static populations’ switch to behavior *b*. A relatively low value of this parameter was enough to cause turnover populations to switch over to *b* faster than static populations. However, it greatly slowed the adoption in all populations, whereas in the experiment, we witnessed a rapid adoption.

To address this, we added a social information bias on choice via value-shaping (Najar et al., 2020). This meant that the rewards obtained by neighboring agents influenced the expected values of the focal agent. A social information bias was evidenced in prior analyses of great tit behavior (Aplin et al., 2017). Further, if a species can socially learn novel behavior, they should also be generally sensitive to social information. The introduction of value shaping reduced the time it took turnover populations to adopt behavior *b*, generating data very similar to our experimental data (shown in figure A1). Social influence also caused a sharp increase the the relative valuation of the high-reward behavior by polymorphic individuals, which matched our verbal hypothesis that naive learners are better samplers. This resulted in strong benefits of even a small amount of turnover during the initial portion of the simulation (figure A2).

## Discussion

We found that turnover interacted with guided variation during maintenance (Boyd and Richerson, 1988; Galef Jr, 1995) to shape how the distributions of cultural traits evolved. The key mechanism by which turnover operated on culture was by changing the composition of repertoires and thus the expected valuations of behavior. Under optimal regimes of turnover, the relative proportion of high-reward behavior produced by the population increased over the long term, compared to static populations. Here, naive agents were more likely to acquire the high-reward behavior only, since transmission was conditioned on production. However, turnover was a double-edged sword. It decreased the proportion of agents that knew both behaviors, and higher intensity turnover regimes led to frequent behavioral extinction. Strong turnover lead a change in composition of the population from polymorphic, where a majority of individuals knew both behaviors, to monomorphic, where individuals only knew one behavior. This fragmentation of the population appeared similar to information gerrymandering (Stewart et al., 2019), although resulted from the social process of replacement rather than assortativity, and resulted in a diminished carrying capacity for cultural traits. The precise points of transition between increasing and decreasing cultural efficiency depended on properties of the network and learners. Finally, as found in our experiment, turnover could greatly benefit populations in the short term, but only under the assumption of choice autocorrelation and social influence.

Our findings provide theoretical support for the hypothesis that introducing naive individuals can drive the cultural evolution of efficiency simply by being “better” samplers, overcoming the cultural inertia of knowledgeable individuals (Chimento et al., 2021; Warner, 1988, 1990). This is a generalizable prediction, as our model did not assume any particular social learning mechanism, and could represent anything from enhancement to imitation (Hoppitt and Laland, 2013). Importantly, this does not oppose the hypothesis that naive incomers introduce innovations(Sasaki and Biro, 2017), and both mechanisms might operate depending on the behavioral context. Additionally, our hypothesis complements, rather than contrasts, the hypothesis that informational cognitive biases of naive learners, such as a bias for simplicity, shape the evolution of cultural traits, studied in the context of language (Culbertson and Kirby, 2016; Culbertson and Smolensky, 2012; Kirby et al., 2008), music (Ravignani et al., 2018; Savage et al., 2015; Verhoef et al., 2014) the evolution of technology (Thompson and Griffiths, 2021). We did not implement any type of cognitive bias towards one behavior or the other, yet one could imagine that the expression of any latent informational bias would be amplified in the absence of preferences developed through reward or punishment.

Guided variation has been formally modeled before by Boyd and Richerson (Boyd and Rich-erson, 1988) where, similar to our model, individual learning was paired with cultural transmission. Beyond this general similarity, the aims, implementation and conclusions differed from our own. Their model aimed to explore the effect of guided variation over evolutionary time-scales. Complete turnover of the population occurred only after each generations’ lifetime, and each new generation inherited their initial phenotype from the previous generation. Through guided variation, the distribution of initial phenotypes was shifted in a direction best fit to the local environment with some error, becoming a distribution of mature phenotypes. Variants favored by individual learning would become more frequent, at the expense of less favored variants, and thus the initial phenotype of the following generation would be different. Thus, the equilibrium trait frequencies would come to match the optimal value given the environment (pp. 98 (Boyd and Richerson, 1988)), and they concluded that guided variation only hastened the evolution towards equilibrium frequencies. In comparison, static populations in our model obtained a equilibrium point of behavioral frequencies that was conditional on both production parameters and payoff probabilities. We showed that the introduction of naive learners via turnover could change the distribution of behaviors, potentially amplifying the the relative proportion of highreward behaviors beyond the equilibrium point. Boyd and Richerson’s model could not account for changes to population structure that could occur within a generation, and thus missed potentially important consequences of population turnover for guided variation. We therefore suggest that turnover can amplify (or nullify if too strong) the effect of guided variation on cultural change.

Unlike Boyd and Richerson’s model, our agents did not inherit knowledge from replaced individuals, and fitness levels did not influence probabilities of replacement. We therefore suggest that our model’s predictions be interpreted as applying to intra-generational time-scales of animals who exhibit fission-fusion social systems, such as passerine birds (Farine et al., 2015). Not only can fission-fusion benefit the initial diffusion of a novel behavior (Aplin et al., 2015; Nunn et al., 2009), but we have demonstrated how it can result in cultural selection. We did not consider migration between multiple populations in the current model (as in Somveille et al. 2018 (Somveille et al., 2018)), yet it is predictable that the risk of cultural extinction might be mitigated by population mixing via fission-fusion. Movement between populations would distribute behavioral information over larger portions of the population, buffering against acute cultural extinction. This decentralization of repertoire merits further exploration.

However, our results do allow us to speculate on the effects of turnover at evolutionary time-scales. The capacity for horizontal cultural transmission is an adaptive strategy in environments that vary over larger time-scales relative to generation times (i.e. red noise environments)(Whitehead, 2007; Whitehead and Richerson, 2009). Population turnover could therefore be most beneficial for cultural species in the relatively stable periods in-between large environmental changes. Interestingly, fission-fusion dynamics have also been predicted to evolve in such cases of “white noise” variability, when there is predictable temporal variation paired with intermediate levels of spatial variation (Sueur et al., 2011). The loss of polymorphic individuals under strong turnover regimes could lead to traps at longer time-scales, especially if behavior is costly to acquire (either socially or asocially) or if there is no long-term retention of information, e.g., via vertical transmission. In such a case, populations could be composed of monomorphic individuals who are best fit the previous environmental state. Despite this, drastic turnover can still improve fitness when the reinvention of behavior is cheap and obligatory, as illustrated by the example of naive blue-headed wrasse choosing more optimal spawning sites, overcoming cultural inertia (Warner, 1988, 1990). While we did not directly test the performance of populations in variable environments, this would be a fruitful future direction for the model.

Stable social groups and longer memories have been hypothesized to be more adaptive in variable environments, as they allow for the accumulation of rare but important knowledge, such as in matriarch elephants (McComb et al., 2001), and give ample opportunity for the social transmission of complex behaviors (Gruber et al., 2022). However, such species are also often characterized by behavioral conservatism (Davis et al., 2016; Hopper et al., 2011; Hrubesch et al., 2009; Marshall-Pescini and Whiten, 2008; Whitehead et al., 2004), and may make maladaptive, conserved decisions, especially when innovation is costly (Keith and Bull, 2017; Whitehead and Richerson, 2009). This conservatism may be detrimental to the diffusion of novel behavior, and even if some individuals possess a large repertoire of behavioral knowledge, the realized usage of such knowledge might be constrained by conservative decision-making rules (Chimento et al., 2022). We found that when behavioral conservatism was implemented as a risk-appetite bias, turnover didn’t benefit risk-averse populations as much as risk-neutral populations. Yet when we implemented it as a choice autocorrelation parameter, performance drastically improved in the short term. This suggests there is nuance to the hypothesis that population turnover is a key mechanism of cultural evolution in behaviorally conservative species (Chimento et al., 2021), in that it might depend on the cognitive mechanism causing conservatism.

## Conclusion

In summary, our study has provided a more thorough understanding of how a social process can influence the internal states of individuals, both in terms of repertoire and valuation. Turnover provided the opportunity for a population to change it’s composition. The precise details of the re-composition process depended on the network size and structure, transmissibility of behaviors, and the relative competitiveness between behaviors, controlled by learning parameters. Intra-generational turnover is commonly found in many species via fission-fusion dynamics and dispersal (Silk et al., 2014), and the introduction of naive learners is a common occurrence in such a social system. Our results suggest that turnover is an important mechanism of cultural evolution, as much so as the better studied social factors of population size (Henrich, 2004) or network connectivity (Derex and Boyd, 2016). Optimal turnover regimes could lead to cultural change without transmission biases or meta-cognition about the choice of demonstrators (Caldwell et al., 2020; Heyes, 2016, 2018; Laland, 2004). However turnover can also constrain the carrying capacity for culture, and lead to extirpation. We demonstrated these beneficial and detrimental effects of turnover with the minimal assumptions that 1) agents could learn from reward, and that 2) the transmission of a behavior depended on the number of associates and the frequencies of their productions. Thus, we suggest the prediction that the introduction of naive learners can itself drive cultural selection is broadly applicable across socially learning species. We emphasize that our model framework, introduced in Chimento et al. (2022), allows researchers to think through factors of transmission and reinforcement learning that might influence culture, implement them and explore their consequences.

## Acknowledgments

MC thanks Wataru Toyokawa for his insightful comments on the initial draft of this manuscript. MC received funding from the IMPRS-QBEE and the Centre for the Advanced Study of Collective Behaviour, funded by the Deutsche Forschungsgemeinschaft (DFG) under Germany’s Excellence Strategy (EXC 2117-422037984). LMA was funded by a Max Planck Research Group Leader Fellowship from the Max Planck Society. This work was additionally supported by the Swiss State Secretariat for Education, Research and Innovation (SERI) under contract number MB22.00056.

## Statement of Authorship

Conceptualisation, M.C., L.M.A.; Methodology, M.C.; Software, M.C.; Investigation, M.C.; Resources, M.C.; Writing Original Draft, M.C.; Writing Review & Editing, M.C., L.M.A.; Visualization, M.C.; Supervision, L.M.A.; Funding Acquisition, L.M.A.

## Data and Code Availability

Code and data for statistical analyses and main text figures, as well as code to replicate the agent based model will be made publicly available upon acceptance. For peer review, the data and code are available for download at https://owncloud.gwdg.de/index.php/s/jgS113lj3CtwBFD.

## Appendix A

### Additional parameterizations

#### 0.1 Modifications required to simulate experimental dynamics

In the main text, we simplified the model as much as possible. However, the flexibility of our model allows for nearly limitless parameterizations of either sub-model. We detail the changes made to the model to better capture the dynamics found in our prior experiment with great tits. First, great tits in static populations rarely sampled the high-reward alternative, and it only became the preferred the solution in 1 of 9 static populations. In contrast, agents in our model adopted *b* quite quickly, usually before turnover populations did. We introduced a simple choice autocorrelation factor by modifying Equation 6 to the following:

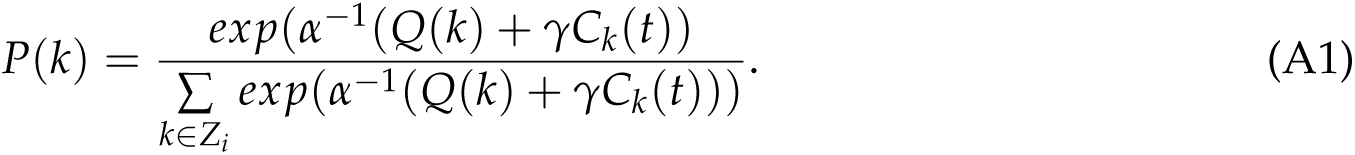

Here, *C_k_*(*t*) records how frequently the focal individual produced behavior *k* within slidingwindow *m*. Parameter *γ* controls the tendency of avoiding or repeating recently chosen behaviors, depending on whether it is negative or positive. We set *γ* = .2. We set *m* = 10, and note that larger *m* values result in a stronger effect of *γ* at a given value.

Second, the adoption curve of the high-reward behavior in turnover populations of great tits was quite steep. Prior modeling work suggested that rapid shifts in preferences could be caused by social influence (Aplin et al., 2017). It seems reasonable that a species which socially learns novel behavior is also subject to social influence on choice. We modified the sub-model of production to include value-shaping, introducing social influence on choices (Najar et al., 2020). Agents observed the rewards that neighboring agents received for producing behaviors and updated their expected values with these rewards, as if they were their own choices:

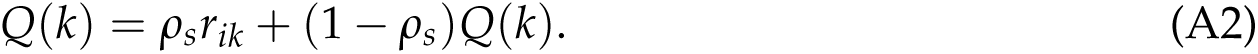

Here, *ρ_s_* was a social learning rate and *r_ik_* was the reward that neighbor *i* received for behavior *k*. We set *ρ_s_* = *ρ* = 0.1 for simplicity, so that socially observed rewards were weighted the same as individually received rewards.

Figure A1 can be compared with Figure 2 to illustrate how the introduction of these two parameters changes dynamics. The diffusion and adoption of behavior *b* is slower in static populations relative to turnover populations, reaching a stable mean proportion of .998 of all behaviors at 824 timesteps in static populations, which was approximately the same in turnover populations at reference level (.997). Given enough time, static and turnover populations would perform equivalently, which is what we expected from the experimental data. Importantly, turnover was immediately beneficial, which we did not find strong evidence for in the simpler parameterization of behavioral conservatism in the main text. The relative valuation of behaviors in turnover populations rose sharply at the beginning of the simulation. This matched our verbal hypothesis as to why turnover matters more for behaviorally conservative species, since naive learners are ”better” samplers of the behavior space than conservative, long-term residents. When we zoom in to the first 824 timesteps (the average time to full diffusion in static condition), we see that turnover could lead to over 30% increases in relative efficiency in figure A2.

---

**Figure A1:**
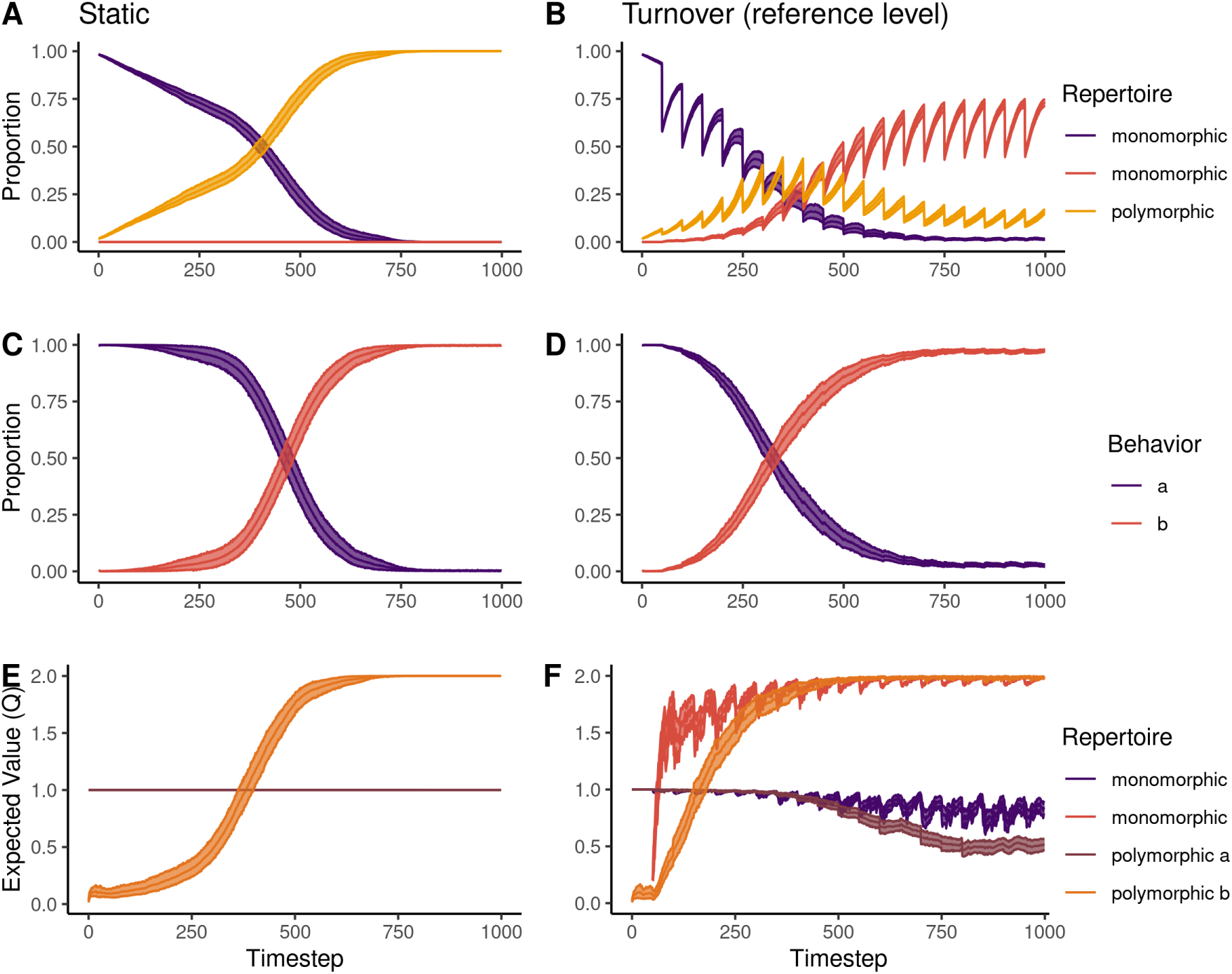
Temporal dynamics under behavioral autocorrelation and social influence. Mean and bootstrapped 95% CL of 3 key metrics (rows) in static and turnover reference level simulations over the first 1000 timesteps. The addition of choice autocorrelation and social influence, generated dynamics more similar to the results and verbal hypothesis from our original experiment with great tits. Turnover populations switch over to behavior *b* faster than static populations, and have a greatly elevated valuation of behavior *b* from the beginning of the simulation.

**Figure A2:**
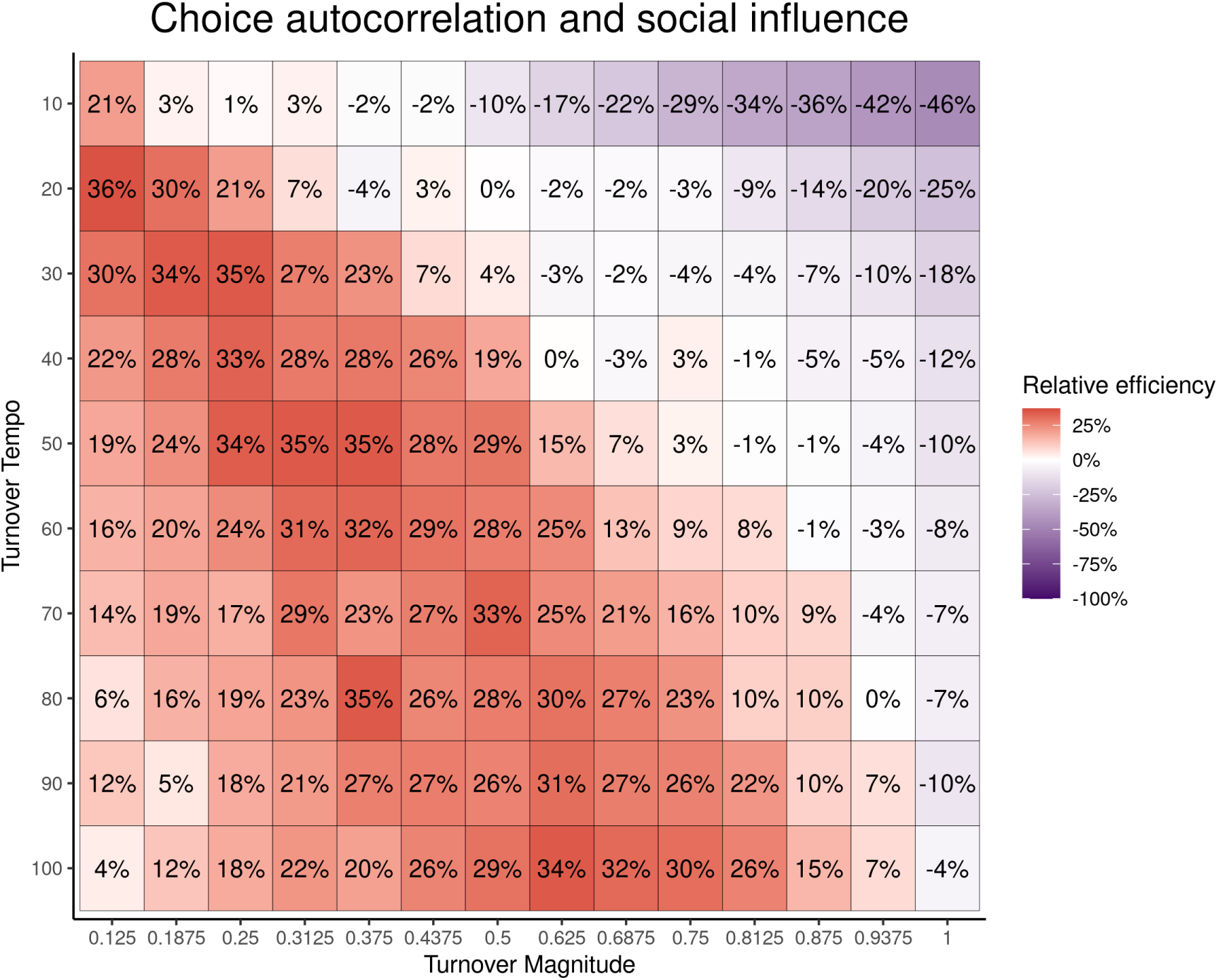
Relative efficiency when populations have autocorrelated choice and social influence. Mean relative efficiency (fill, label) for 100 simulations under each turnover regime: the combination of turnover magnitude (x-axis) and turnover tempo (y-axis). Relative efficiency was measured against the proportion of *b* produced by the static populations under a given condition, where the color white is equivalent performance, salmon is over-performance and purple is under-performance. Data is subset to during the initial 824 timesteps of simulations to highlight how even small amounts of turnover yield short term benefits for behaviorally conservative spopulations.

## Literature Cited

1. Acerbi, A., and R. A. Bentley. 2014. Biases in cultural transmission shape the turnover of popular traits. Evolution and Human Behavior 35:228–236.

2. Aplin, L. M. 2019. Culture and cultural evolution in birds: a review of the evidence. Animal Behaviour 147:179–187.

3. Aplin, L. M., D. R. Farine, J. Morand-Ferron, A. Cockburn, A. Thornton, and B. C. Sheldon. 2015. Experimentally induced innovations lead to persistent culture via conformity in wild birds. Nature 518:538.

4. Aplin, L. M., B. C. Sheldon, and R. McElreath. 2017. Conformity does not perpetuate suboptimal traditions in a wild population of songbirds. Proceedings of the National Academy of Sciences 114:7830–7837.

5. Aureli, F., C. M. Schaffner, C. Boesch, S. K. Bearder, J. Call, C. A. Chapman, R. Connor, A. D. Fiore, R. I. Dunbar, S. P. Henzi, et al. 2008. Fission-fusion dynamics: new research frameworks. Current Anthropology 49:627–654.

6. Bass, F. M. 1969. A new product growth for model consumer durables. Management science 15:215–227.

7. Bentley, R. A., C. P. Lipo, H. A. Herzog, and M. W. Hahn. 2007. Regular rates of popular culture change reflect random copying. Evolution and Human Behavior 28:151–158.

8. Boyd, R., and P. J. Richerson. 1988. Culture and the evolutionary process. University of Chicago press.

9. Caldwell, C. A., M. Atkinson, K. H. Blakey, J. Dunstone, D. Kean, G. Mackintosh, E. Renner, and C. E. Wilks. 2020. Experimental assessment of capacities for cumulative culture: Review and evaluation of methods. Wiley Interdisciplinary Reviews: Cognitive Science 11:e1516.

10. Chimento, M., G. Alarcón-Nieto, and L. M. Aplin. 2021. Population turnover facilitates cultural selection for efficiency in birds. Current Biology 31:2477–2483.

11. Chimento, M., B. J. Barrett, A. Kandler, and L. M. Aplin. 2022. Cultural diffusion dynamics depend on behavioural production rules. Proceedings of the Royal Society B 289:20221001.

12. Creanza, N., O. Kolodny, and M. W. Feldman. 2017a. Cultural evolutionary theory: How culture evolves and why it matters. Proceedings of the National Academy of Sciences 114:7782–7789.

13. Creanza, N., O. Kolodny, and M. W. Feldman. 2017b. Greater than the sum of its parts? modelling population contact and interaction of cultural repertoires. Journal of The Royal Society Interface 14:20170171.

14. Culbertson, J., and S. Kirby. 2016. Simplicity and specificity in language: Domain-general biases have domain-specific effects. Frontiers in psychology 6:1964.

15. Culbertson, J., and P. Smolensky. 2012. A bayesian model of biases in artificial language learning: The case of a word-order universal. Cognitive science 36:1468–1498.

16. Dale, R., and G. Lupyan. 2012. Understanding the origins of morphological diversity: The linguistic niche hypothesis. Advances in complex systems 15:1150017.

17. Davis, S. J., G. L. Vale, S. J. Schapiro, S. P. Lambeth, and A. Whiten. 2016. Foundations of cumulative culture in apes: improved foraging efficiency through relinquishing and combining witnessed behaviours in chimpanzees (pan troglodytes). Scientific reports 6:35953.

18. Dean, L. G., R. L. Kendal, S. J. Schapiro, B. Thierry, and K. N. Laland. 2012. Identification of the social and cognitive processes underlying human cumulative culture. Science 335:1114–1118.

19. Deffner, D., and R. McElreath. 2022. When does selection favor learning from the old? social learning in age-structured populations. PloS one 17:e0267204.

20. Derex, M., and R. Boyd. 2016. Partial connectivity increases cultural accumulation within groups. Proceedings of the National Academy of Sciences 113:2982–2987.

21. Farine, D. R., J. A. Firth, L. M. Aplin, R. A. Crates, A. Culina, C. J. Garroway, C. A. Hinde, L. R. Kidd, N. D. Milligan, I. Psorakis, et al. 2015. The role of social and ecological processes in structuring animal populations: a case study from automated tracking of wild birds. Royal Society Open Science 2:150057.

22. Franz, M., and C. L. Nunn. 2009. Network-based diffusion analysis: a new method for detecting social learning. Proceedings of the Royal Society B: Biological Sciences 276:1829–1836.

23. Galef Jr, B. G. 1995. Why behaviour patterns that animals learn socially are locally adaptive. Animal behaviour 49:1325–1334.

24. Gruber, T., M. Chimento, L. M. Aplin, and D. Biro. 2022. Efficiency fosters cumulative culture across species. Philosophical Transactions of the Royal Society B 377:20200308.

25. Hasenjager, M. J., E. Leadbeater, and W. Hoppitt. 2020. Detecting and quantifying social transmission using network-based diffusion analysis. Journal of Animal Ecology .

26. Henrich, J. 2004. Demography and cultural evolution: how adaptive cultural processes can produce maladaptive losses—the tasmanian case. American antiquity 69:197–214.

27. Heyes, C. 2016. Who knows? metacognitive social learning strategies. Trends in cognitive sciences 20:204–213.

28. Heyes, C. 2018. Enquire within: Cultural evolution and cognitive science. Philosophical Transactions of the Royal Society B: Biological Sciences 373:20170051.

29. Hopper, L. M., S. J. Schapiro, S. P. Lambeth, and S. F. Brosnan. 2011. Chimpanzees’ socially maintained food preferences indicate both conservatism and conformity. Animal Behaviour 81:1195–1202.

30. Hoppitt, W., and K. N. Laland. 2013. Social learning: an introduction to mechanisms, methods, and models. Princeton University Press.

31. Hrubesch, C., S. Preuschoft, and C. van Schaik. 2009. Skill mastery inhibits adoption of observed alternative solutions among chimpanzees (pan troglodytes). Animal cognition 12:209.

32. Katahira, K. 2015. The relation between reinforcement learning parameters and the influence of reinforcement history on choice behavior. Journal of Mathematical Psychology 66:59–69.

33. Keith, S. A., and J. W. Bull. 2017. Animal culture impacts species’ capacity to realise climate-driven range shifts. Ecography 40:296–304.

34. Kendal, R. L., N. J. Boogert, L. Rendell, K. N. Laland, M. Webster, and P. L. Jones. 2018. Social learning strategies: Bridge-building between fields. Trends in cognitive sciences 22:651–665.

35. Kirby, S., H. Cornish, and K. Smith. 2008. Cumulative cultural evolution in the laboratory: An experimental approach to the origins of structure in human language. Proceedings of the National Academy of Sciences 105:10681–10686.

36. Kirby, S., T. Griffiths, and K. Smith. 2014. Iterated learning and the evolution of language. Current opinion in neurobiology 28:108–114.

37. Kirby, S., and M. Tamariz. 2022. Cumulative cultural evolution, population structure and the origin of combinatoriality in human language. Philosophical Transactions of the Royal Society B 377:20200319.

38. Laland, K. N. 2004. Social learning strategies. Animal Learning & Behavior 32:4–14.

39. Legare, C. H. 2017. Cumulative cultural learning: Development and diversity. Proceedings of the National Academy of Sciences 114:7877–7883.

40. Lupyan, G., and R. Dale. 2010. Language structure is partly determined by social structure. PloS one 5:e8559.

41. Marshall-Pescini, S., and A. Whiten. 2008. Chimpanzees (pan troglodytes) and the question of cumulative culture: an experimental approach. Animal cognition 11:449–456.

42. McComb, K., C. Moss, S. M. Durant, L. Baker, and S. Sayialel. 2001. Matriarchs as repositories of social knowledge in african elephants. Science 292:491–494.

43. Mesoudi, A. 2017. Pursuing darwin’s curious parallel: Prospects for a science of cultural evolution. Proceedings of the National Academy of Sciences 114:7853–7860.

44. Mesoudi, A., and A. Thornton. 2018. What is cumulative cultural evolution? Proceedings of the Royal Society B: Biological Sciences 285:20180712.

45. Mesoudi, A., and A. Whiten. 2008. The multiple roles of cultural transmission experiments in understanding human cultural evolution. Philosophical Transactions of the Royal Society B: Biological Sciences 363:3489–3501.

46. Mesoudi, A., A. Whiten, and K. N. Laland. 2004. Perspective: Is human cultural evolution darwinian? evidence reviewed from the perspective of the origin of species. Evolution 58:1–11.

47. Najar, A., E. Bonnet, B. Bahrami, and S. Palminteri. 2020. The actions of others act as a pseudo-reward to drive imitation in the context of social reinforcement learning. PLoS biology 18:e3001028.

48. Nunn, C. L., P. H. Thrall, K. Bartz, T. Dasgupta, and C. Boesch. 2009. Do transmission mechanisms or social systems drive cultural dynamics in socially structured populations? Animal Behaviour 77:1515–1524.

49. Ravignani, A., B. Thompson, T. Grossi, T. Delgado, and S. Kirby. 2018. Evolving building blocks of rhythm: how human cognition creates music via cultural transmission. Annals of the New York Academy of Sciences 1423:176–187.

50. Raviv, L., A. Meyer, and S. Lev-Ari. 2019. Larger communities create more systematic languages. Proceedings of the Royal Society B 286:20191262.

51. Rendell, L., L. Fogarty, W. J. Hoppitt, T. J. Morgan, M. M. Webster, and K. N. Laland. 2011. Cognitive culture: theoretical and empirical insights into social learning strategies. Trends in cognitive sciences 15:68–76.

52. Rendell, L., and H. Whitehead. 2001. Culture in whales and dolphins. Behavioral and brain sciences 24:309–324.

53. Sasaki, T., and D. Biro. 2017. Cumulative culture can emerge from collective intelligence in animal groups. Nature communications 8:1–6.

54. Savage, P. E., S. Brown, E. Sakai, and T. E. Currie. 2015. Statistical universals reveal the structures and functions of human music. Proceedings of the National Academy of Sciences 112:8987– 8992.

55. Schuppli, C., and C. P. van Schaik. 2019. Animal cultures: How we’ve only seen the tip of the iceberg. Evolutionary Human Sciences 1.

56. Silk, M. J., D. P. Croft, T. Tregenza, and S. Bearhop. 2014. The importance of fission–fusion social group dynamics in birds. Ibis 156:701–715.

57. Smith, K. 2002. The cultural evolution of communication in a population of neural networks. Connection Science 14:65–84.

58. Somveille, M., J. A. Firth, L. M. Aplin, D. R. Farine, B. C. Sheldon, and R. N. Thompson. 2018. Movement and conformity interact to establish local behavioural traditions in animal populations. PLoS computational biology 14:e1006647.

59. Stewart, A. J., M. Mosleh, M. Diakonova, A. A. Arechar, D. G. Rand, and J. B. Plotkin. 2019. Information gerrymandering and undemocratic decisions. Nature 573:117–121.

60. Sueur, C., A. J. King, L. Conradt, G. Kerth, D. Lusseau, C. Mettke-Hofmann, C. M. Schaffner, L. Williams, D. Zinner, and F. Aureli. 2011. Collective decision-making and fission–fusion dynamics: a conceptual framework. Oikos 120:1608–1617.

61. Sutton, R. S., A. G. Barto, et al. 1998. Introduction to reinforcement learning, vol. 2. MIT press Cambridge.

62. Tennie, C., J. Call, and M. Tomasello. 2009. Ratcheting up the ratchet: on the evolution of cumulative culture. Philosophical Transactions of the Royal Society B: Biological Sciences 364:2405– 2415.

63. Thompson, B., and T. L. Griffiths. 2021. Human biases limit cumulative innovation. Proceedings of the Royal Society B 288:20202752.

64. Verhoef, T., S. Kirby, and B. De Boer. 2014. Emergence of combinatorial structure and economy through iterated learning with continuous acoustic signals. Journal of Phonetics 43:57–68.

65. Warner, R. R. 1988. Traditionality of mating-site preferences in a coral reef fish. Nature 335:719– 721.

66. Warner, R. R. 1990. Resource assessment versus tradition in mating-site determination. The American Naturalist 135:205–217.

67. Whitehead, H. 2007. Learning, climate and the evolution of cultural capacity. Journal of theoretical biology 245:341–350.

68. Whitehead, H., and D. Lusseau. 2012. Animal social networks as substrate for cultural behavioural diversity. Journal of theoretical biology 294:19–28.

69. Whitehead, H., L. Rendell, R. W. Osborne, and B. Wü rsig. 2004. Culture and conservation of non-humans with reference to whales and dolphins: review and new directions. Biological Conservation 120:427–437.

70. Whitehead, H., and P. J. Richerson. 2009. The evolution of conformist social learning can cause population collapse in realistically variable environments. Evolution and Human Behavior 30:261–273.

71. Whiten, A. 2019. Social learning: Peering deeper into ape culture. Current Biology 29:R845–R847.

72. Whiten, A. 2021. The burgeoning reach of animal culture. Science 372.

73. Whiten, A., R. A. Hinde, K. N. Laland, and C. B. Stringer. 2011. Culture evolves.

74. Whiten, A., and A. Mesoudi. 2008. Establishing an experimental science of culture: animal social diffusion experiments. Philosophical Transactions of the Royal Society B: Biological Sciences 363:3477–3488.

75. Wray, A., and G. W. Grace. 2007. The consequences of talking to strangers: Evolutionary corollaries of socio-cultural influences on linguistic form. Lingua 117:543–578.

