## Supplementary figures and tables for "From Data to Theory: Understanding the Role of Naive Learners in Cultural Change"

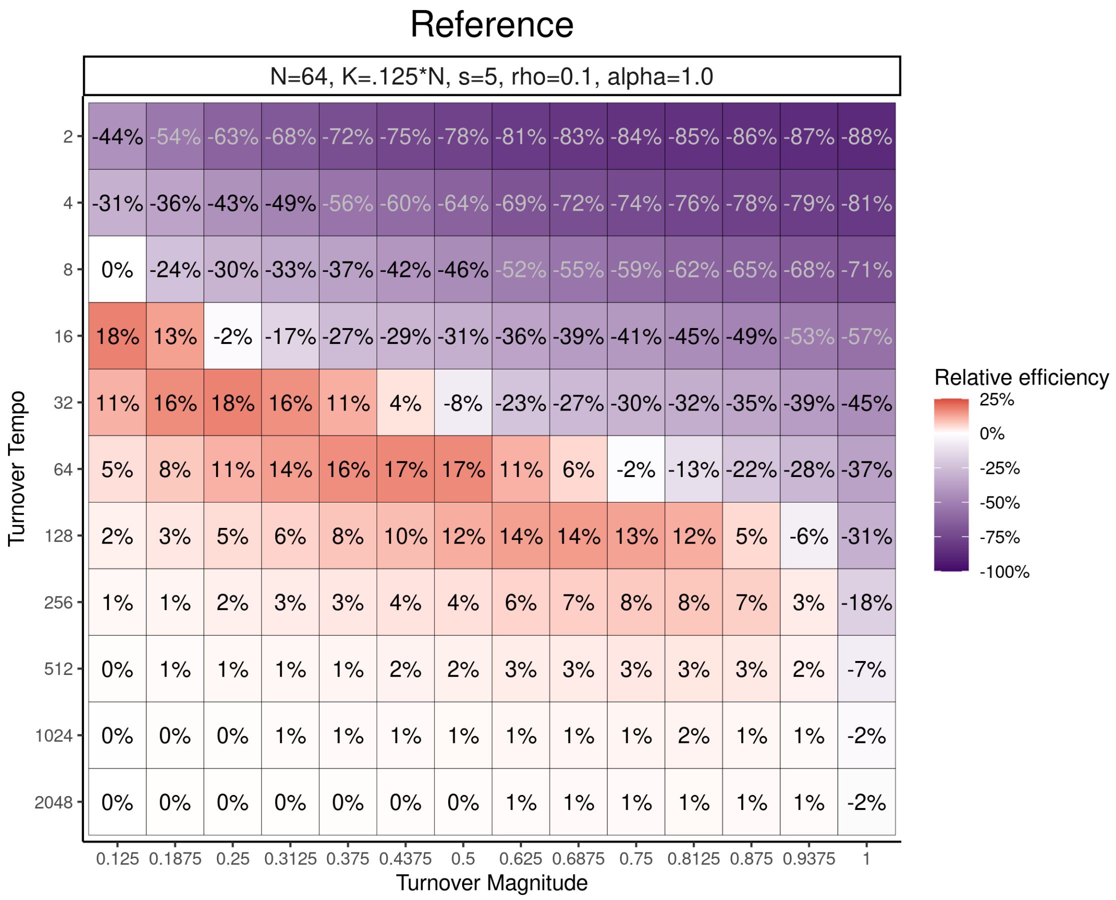


Figure S1: **Extended exploration of turnover tempo.** The effects of turnover magnitude diminished as tempo decreased, and populations were almost no different than static populations at $T_m = 1024$.

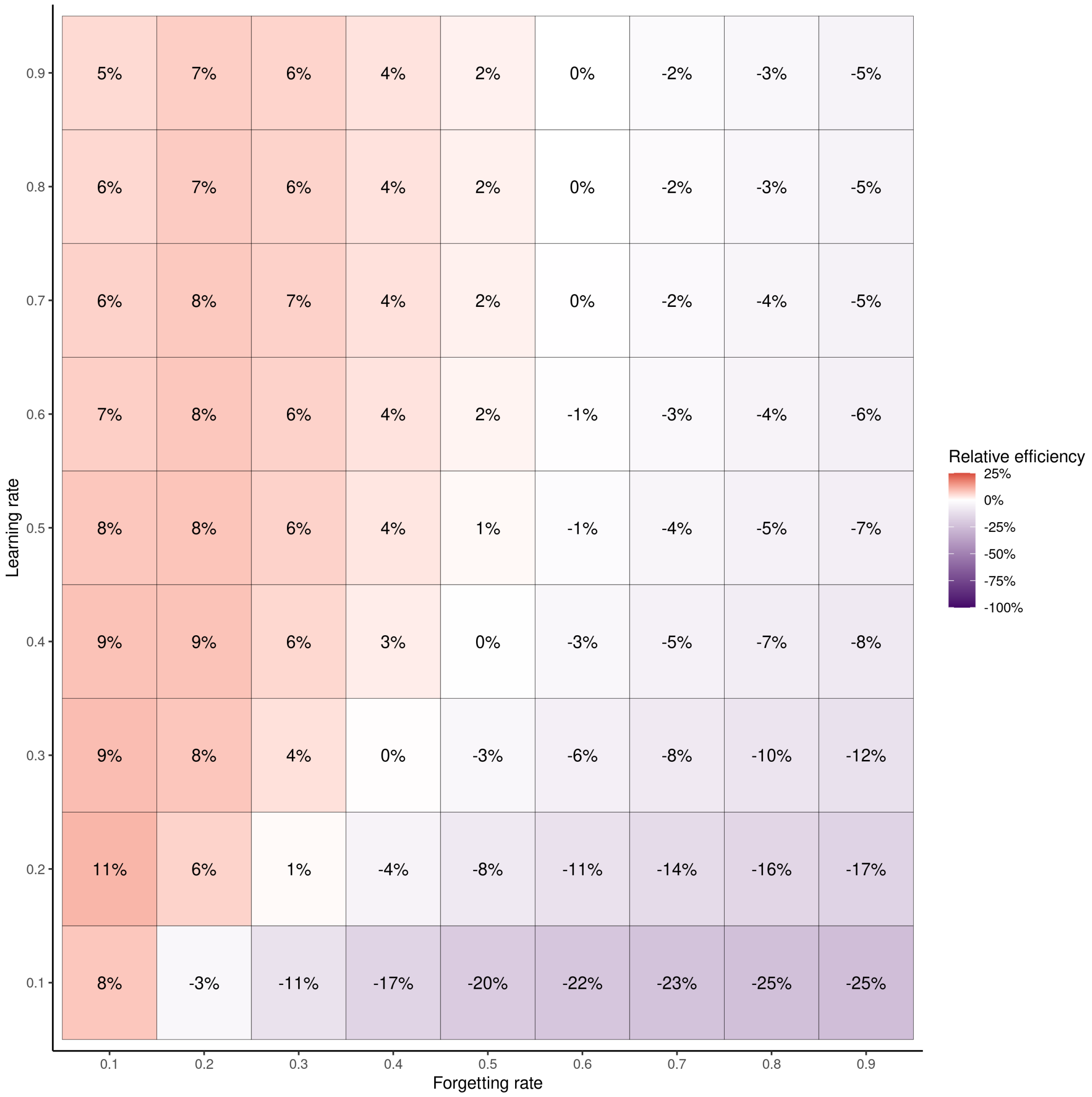


Figure S2: **The effect of forgetting rate on static populations.** Relative efficiency of static populations with varying levels of learning and forgetting rates. Relative efficiency was calculated by comparing the proportion of $b$ produced to static populations where $\rho_f = 0$.

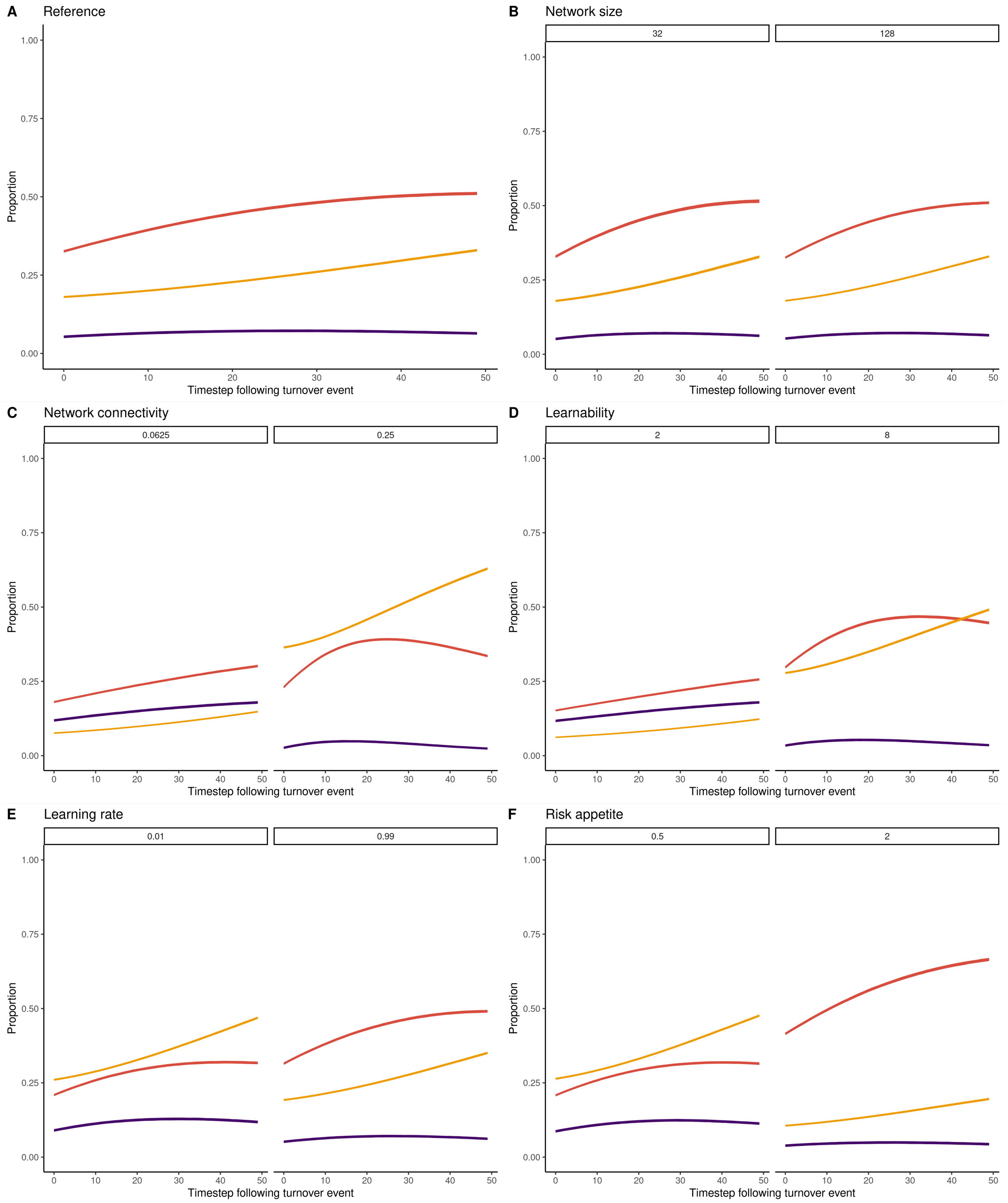


Figure S3: **Population re-composition following turnover event for other parameters.** Mean and 95% CL proportion of agents which were low-reward monomorphic (purple), high-reward monomorphic (red), or polymorphic (yellow). Each panel shows alternative parameter values while all other parameters were held at the reference constellation.

Table S1: **Summary of results for turnover magnitude**. Summary presented for average values taken over all timesteps with optimal magnitude in italics. Reported are the mean and SD values for the proportion of agents knowledgeable about either low-payoff behavior *a* or high-payoff behavior *b* (monomorphic agents) or both behaviors (polymorphic agents), median proportion of behaviors that are either (*a*) or (*b*) within a given timestep, median expected payoffs held by knowledgeable agents for *a* and *b*.

| | low-reward | | high-reward | | both | |
|---|---|---|---|---|---|---|
| magnitude | mean | sd | mean | sd | mean | sd |
| **Repertoire state** | | | | | | |
| static | 0.028 | 0.140 | 0.000 | 0.000 | 0.972 | 0.140 |
| 0.125 | 0.048 | 0.136 | 0.180 | 0.066 | 0.707 | 0.114 |
| 0.1875 | 0.053 | 0.134 | 0.289 | 0.101 | 0.556 | 0.110 |
| 0.25 | 0.058 | 0.134 | 0.379 | 0.130 | 0.419 | 0.111 |
| 0.3125 | 0.062 | 0.133 | 0.431 | 0.146 | 0.318 | 0.108 |
| *0.375* | *0.067* | *0.131* | *0.448* | *0.155* | *0.246* | *0.101* |
| 0.4375 | 0.072 | 0.128 | 0.438 | 0.156 | 0.195 | 0.092 |
| 0.5 | 0.086 | 0.130 | 0.395 | 0.156 | 0.162 | 0.084 |
| 0.625 | 0.116 | 0.130 | 0.282 | 0.143 | 0.101 | 0.064 |
| 0.6875 | 0.126 | 0.129 | 0.217 | 0.130 | 0.070 | 0.053 |
| 0.75 | 0.122 | 0.126 | 0.163 | 0.113 | 0.045 | 0.041 |
| 0.8125 | 0.107 | 0.121 | 0.116 | 0.094 | 0.026 | 0.030 |
| 0.875 | 0.089 | 0.114 | 0.082 | 0.077 | 0.014 | 0.021 |
| 0.9375 | 0.068 | 0.108 | 0.060 | 0.064 | 0.007 | 0.015 |
| 1 | 0.052 | 0.104 | 0.044 | 0.053 | 0.004 | 0.011 |
| **Behavioral productions** | | | | | | |
| static | 0.291 | 0.121 | 0.709 | 0.121 | | |
| 0.125 | 0.247 | 0.132 | 0.753 | 0.132 | | |
| 0.1875 | 0.216 | 0.140 | 0.784 | 0.140 | | |
| 0.25 | 0.189 | 0.150 | 0.811 | 0.150 | | |
| 0.3125 | 0.173 | 0.157 | 0.827 | 0.157 | | |
| *0.375* | *0.169* | *0.163* | *0.831* | *0.163* | | |
| 0.4375 | 0.172 | 0.166 | 0.828 | 0.166 | | |
| 0.5 | 0.198 | 0.179 | 0.802 | 0.179 | | |
| 0.625 | 0.286 | 0.206 | 0.714 | 0.206 | | |
| 0.6875 | 0.350 | 0.227 | 0.650 | 0.227 | | |
| 0.75 | 0.404 | 0.244 | 0.595 | 0.244 | | |
| 0.8125 | 0.448 | 0.266 | 0.551 | 0.266 | | |
| 0.875 | 0.481 | 0.292 | 0.509 | 0.293 | | |
| 0.9375 | 0.473 | 0.333 | 0.481 | 0.333 | | |
| 1 | 0.427 | 0.363 | 0.431 | 0.364 | | |
| **Expected payoffs** | | | | | | |
| static | 0.998 | 0.029 | 1.972 | 0.146 | | |
| 0.125 | 0.880 | 0.045 | 1.909 | 0.141 | | |
| 0.1875 | 0.819 | 0.058 | 1.879 | 0.138 | | |
| 0.25 | 0.768 | 0.073 | 1.848 | 0.139 | | |
| 0.3125 | 0.730 | 0.086 | 1.816 | 0.137 | | |
| *0.375* | *0.705* | *0.098* | *1.782* | *0.138* | | |
| 0.4375 | 0.688 | 0.109 | 1.745 | 0.140 | | |
| 0.5 | 0.686 | 0.116 | 1.703 | 0.142 | | |
| 0.625 | 0.700 | 0.125 | 1.610 | 0.159 | | |
| 0.6875 | 0.710 | 0.129 | 1.562 | 0.181 | | |
| 0.75 | 0.711 | 0.139 | 1.517 | 0.215 | | |
| 0.8125 | 0.707 | 0.153 | 1.471 | 0.266 | | |
| 0.875 | 0.694 | 0.169 | 1.410 | 0.328 | | |
| 0.9375 | 0.661 | 0.192 | 1.327 | 0.376 | | |
| 1 | 0.602 | 0.203 | 1.203 | 0.400 | | |

Table S2: **Summary of results for turnover tempo**. Summary presented for average values taken over all timesteps with optimal tempo in italics. Reported are the mean and SD values for the proportion of agents knowledgeable about either low-payoff behavior *a* or high-payoff behavior *b* (monomorphic agents) or both behaviors (polymorphic agents), median proportion of behaviors that are either (*a*) or (*b*) within a given timestep, median expected payoffs held by knowledgeable agents for *a* and *b*.

| | low-reward | | high-reward | | both | |
|---|---|---|---|---|---|---|
| tempo | mean | sd | mean | sd | mean | sd |
| **Repertoire state** | | | | | | |
| static | 0.028 | 0.140 | 0.000 | 0.000 | 0.972 | 0.140 |
| 100 | 0.054 | 0.134 | 0.290 | 0.107 | 0.543 | 0.133 |
| 90 | 0.056 | 0.135 | 0.325 | 0.117 | 0.491 | 0.128 |
| 80 | 0.057 | 0.135 | 0.362 | 0.128 | 0.438 | 0.124 |
| 70 | 0.058 | 0.133 | 0.402 | 0.139 | 0.374 | 0.117 |
| 60 | 0.062 | 0.133 | 0.433 | 0.150 | 0.310 | 0.111 |
| *50* | *0.067* | *0.131* | *0.448* | *0.155* | *0.246* | *0.101* |
| 40 | 0.076 | 0.126 | 0.431 | 0.150 | 0.185 | 0.084 |
| 30 | 0.099 | 0.122 | 0.348 | 0.137 | 0.120 | 0.063 |
| 20 | 0.123 | 0.110 | 0.162 | 0.097 | 0.040 | 0.032 |
| 10 | 0.038 | 0.064 | 0.034 | 0.032 | 0.002 | 0.006 |
| **Behavioral productions** | | | | | | |
| static | 0.291 | 0.121 | 0.709 | 0.121 | | |
| 100 | 0.216 | 0.139 | 0.784 | 0.139 | | |
| 90 | 0.205 | 0.143 | 0.795 | 0.143 | | |
| 80 | 0.194 | 0.147 | 0.806 | 0.147 | | |
| 70 | 0.181 | 0.151 | 0.819 | 0.151 | | |
| 60 | 0.172 | 0.158 | 0.828 | 0.158 | | |
| *50* | *0.169* | *0.163* | *0.831* | *0.163* | | |
| 40 | 0.177 | 0.169 | 0.823 | 0.169 | | |
| 30 | 0.230 | 0.188 | 0.770 | 0.188 | | |
| 20 | 0.412 | 0.245 | 0.588 | 0.245 | | |
| 10 | 0.475 | 0.353 | 0.477 | 0.353 | | |
| **Expected payoffs** | | | | | | |
| static | 0.998 | 0.029 | 1.972 | 0.146 | | |
| 100 | 0.816 | 0.061 | 1.874 | 0.144 | | |
| 90 | 0.797 | 0.066 | 1.863 | 0.141 | | |
| 80 | 0.776 | 0.072 | 1.851 | 0.141 | | |
| 70 | 0.752 | 0.079 | 1.835 | 0.141 | | |
| 60 | 0.728 | 0.087 | 1.813 | 0.140 | | |
| *50* | *0.705* | *0.098* | *1.782* | *0.138* | | |
| 40 | 0.684 | 0.109 | 1.734 | 0.137 | | |
| 30 | 0.685 | 0.121 | 1.649 | 0.141 | | |
| 20 | 0.709 | 0.129 | 1.502 | 0.205 | | |
| 10 | 0.616 | 0.201 | 1.238 | 0.397 | | |

Table S3: **Summary of expected payoffs for agents in each knowledge state**. Mean and SD expected payoffs for behaviors *a* and *b* held by monomorphic and polymorphic agents in the final 1000 timesteps of simulation. Data presented with all other parameters held at reference setting, optimal levels of magnitude and tempo are italicized. Static populations were composed of only polymorphic agents. At optimal levels of turnover magnitude ($T_m$ = 0.25) and turnover tempo ($T_t$ = 20), polymorphic agents held lower expected payoffs of *a* and higher expected payoffs of *b* compared to monomorphic agents. Their expected payoffs were also exaggerated compared to polymorphic agents in static populations. This was because most agents who became polymorphic had first acquired *b*, followed by *a*. Thus, they accrued more experience with *b* before becoming polymorphic.

| | monomorphic a | | monomorphic b | | polymorphic a | | polymorphic b | |
|---|---|---|---|---|---|---|---|---|
| value | mean | sd | mean | sd | mean | sd | mean | sd |
| **Static** | | | | | | | | |
| static | 0.972 | 0.112 | NA | NA | 0.996 | 0.043 | 1.972 | 0.146 |
| **Turnover magnitude** | | | | | | | | |
| 0.125 | 0.805 | 0.217 | 1.822 | 0.147 | 0.881 | 0.051 | 1.935 | 0.145 |
| 0.1875 | 0.801 | 0.207 | 1.822 | 0.133 | 0.818 | 0.063 | 1.915 | 0.144 |
| 0.25 | 0.798 | 0.200 | 1.814 | 0.130 | 0.765 | 0.077 | 1.89 | 0.149 |
| 0.3125 | 0.795 | 0.195 | 1.796 | 0.127 | 0.722 | 0.091 | 1.859 | 0.154 |
| *0.375* | *0.794* | *0.187* | *1.772* | *0.128* | *0.69* | *0.106* | *1.823* | *0.166* |
| 0.4375 | 0.792 | 0.179 | 1.743 | 0.130 | 0.663 | 0.122 | 1.781 | 0.184 |
| 0.5 | 0.792 | 0.168 | 1.706 | 0.139 | 0.65 | 0.137 | 1.731 | 0.208 |
| 0.625 | 0.784 | 0.147 | 1.625 | 0.165 | 0.634 | 0.177 | 1.609 | 0.288 |
| 0.6875 | 0.776 | 0.140 | 1.581 | 0.194 | 0.623 | 0.208 | 1.546 | 0.356 |
| 0.75 | 0.763 | 0.142 | 1.537 | 0.230 | 0.611 | 0.245 | 1.489 | 0.425 |
| 0.8125 | 0.744 | 0.153 | 1.487 | 0.279 | 0.593 | 0.276 | 1.436 | 0.493 |
| 0.875 | 0.717 | 0.170 | 1.427 | 0.335 | 0.577 | 0.302 | 1.358 | 0.554 |
| 0.9375 | 0.678 | 0.191 | 1.341 | 0.379 | 0.548 | 0.317 | 1.271 | 0.594 |
| 1 | 0.615 | 0.205 | 1.222 | 0.404 | 0.476 | 0.316 | 1.109 | 0.599 |
| **Turnover tempo** | | | | | | | | |
| 100 | 0.808 | 0.211 | 1.823 | 0.163 | 0.816 | 0.064 | 1.912 | 0.146 |
| 90 | 0.807 | 0.207 | 1.819 | 0.155 | 0.796 | 0.070 | 1.904 | 0.145 |
| 80 | 0.799 | 0.206 | 1.815 | 0.146 | 0.774 | 0.075 | 1.894 | 0.146 |
| 70 | 0.798 | 0.202 | 1.807 | 0.140 | 0.748 | 0.082 | 1.88 | 0.150 |
| 60 | 0.794 | 0.197 | 1.794 | 0.134 | 0.72 | 0.092 | 1.858 | 0.156 |
| *50* | *0.794* | *0.187* | *1.772* | *0.128* | *0.69* | *0.106* | *1.823* | *0.166* |
| 40 | 0.791 | 0.174 | 1.732 | 0.128 | 0.656 | 0.124 | 1.766 | 0.189 |
| 30 | 0.786 | 0.153 | 1.66 | 0.136 | 0.628 | 0.155 | 1.653 | 0.250 |
| 20 | 0.759 | 0.130 | 1.526 | 0.211 | 0.594 | 0.248 | 1.453 | 0.440 |
| 10 | 0.625 | 0.199 | 1.248 | 0.396 | 0.48 | 0.318 | 1.128 | 0.618 |

Table S4: **Summary of results for other parameters**. Summary presented for the final 5000 time-steps with tempo and magnitude held at reference level ($T_t$ = 50, $T_m$ = .5). Reported are the median $[Q1, Q3]$ values for the proportion of agents knowledgeable about either low-payoff behavior *a* or high-payoff behavior *b* (monomorphic agents) or both behaviors (polymorphic agents), median proportion of behaviors that are either (*a*) or (*b*) within a given timestep, median expected payoffs held by knowledgeable agents for *a* and *b*.

| | | low-reward | | high-reward | | both | |
|---|---|---|---|---|---|---|---|
| parameter | value | mean | sd | mean | sd | mean | sd |
| **Repertoire state** | | | | | | | |
| Network size | 32.000 | 0.066 | 0.131 | 0.452 | 0.181 | 0.245 | 0.125 |
| Network size | 128.000 | 0.067 | 0.131 | 0.447 | 0.138 | 0.246 | 0.084 |
| Network connectivity | 0.062 | 0.153 | 0.130 | 0.246 | 0.106 | 0.107 | 0.050 |
| Network connectivity | 0.250 | 0.039 | 0.111 | 0.354 | 0.121 | 0.489 | 0.131 |
| Transmissibility | 2.000 | 0.151 | 0.127 | 0.207 | 0.096 | 0.088 | 0.045 |
| Transmissibility | 8.000 | 0.046 | 0.122 | 0.429 | 0.144 | 0.376 | 0.124 |
| Rho | 0.010 | 0.119 | 0.135 | 0.289 | 0.127 | 0.353 | 0.107 |
| Rho | 0.990 | 0.066 | 0.122 | 0.432 | 0.143 | 0.262 | 0.099 |
| Alpha | 0.500 | 0.115 | 0.130 | 0.289 | 0.118 | 0.358 | 0.107 |
| Alpha | 2.000 | 0.046 | 0.133 | 0.569 | 0.172 | 0.146 | 0.086 |
| **Behavioral productions** | | | | | | | |
| Network size | 32.000 | 0.167 | 0.173 | 0.833 | 0.173 | | |
| Network size | 128.000 | 0.168 | 0.157 | 0.832 | 0.157 | | |
| Network connectivity | 0.062 | 0.354 | 0.187 | 0.646 | 0.187 | | |
| Network connectivity | 0.250 | 0.177 | 0.126 | 0.823 | 0.126 | | |
| Transmissibility | 2.000 | 0.384 | 0.192 | 0.616 | 0.192 | | |
| Transmissibility | 8.000 | 0.161 | 0.142 | 0.839 | 0.142 | | |
| Rho | 0.010 | 0.332 | 0.165 | 0.668 | 0.165 | | |
| Rho | 0.990 | 0.176 | 0.146 | 0.824 | 0.146 | | |
| Alpha | 0.500 | 0.328 | 0.150 | 0.672 | 0.150 | | |
| Alpha | 2.000 | 0.083 | 0.169 | 0.917 | 0.169 | | |
| **Expected payoffs** | | | | | | | |
| Network size | 32.000 | 0.700 | 0.139 | 1.784 | 0.148 | | |
| Network size | 128.000 | 0.705 | 0.077 | 1.780 | 0.137 | | |
| Network connectivity | 0.062 | 0.816 | 0.088 | 1.757 | 0.145 | | |
| Network connectivity | 0.250 | 0.669 | 0.077 | 1.791 | 0.151 | | |
| Transmissibility | 2.000 | 0.825 | 0.089 | 1.753 | 0.153 | | |
| Transmissibility | 8.000 | 0.674 | 0.085 | 1.790 | 0.148 | | |
| Rho | 0.010 | 0.290 | 0.059 | 0.750 | 0.129 | | |
| Rho | 0.990 | 0.953 | 0.053 | 1.987 | 0.061 | | |
| Alpha | 0.500 | 0.818 | 0.061 | 1.758 | 0.131 | | |
| Alpha | 2.000 | 0.436 | 0.179 | 1.784 | 0.162 | | |
